# Multi-omic cardiac adaptation to exercise training identifies adaptive and disease-opposing signatures: findings from MoTrPAC

**DOI:** 10.64898/2026.11.16.767197

**Authors:** Pauline Brochet, Gina M. Many, Malene E. Lindholm, Gregory R. Smith, Sasha Gladkikh, Hasmik Keshishian, Daniel Seung Kim, David Jimenez-Morales, Samuel Montalvo, Joyce Njoroge, Sue C. Bodine, Euan A. Ashley, William E. Kraus, Martin J. Walsh, Matthew T. Wheeler, Daniel H. Katz, Writing Group for the MoTrPAC Study Group

## Abstract

**Background:** Endurance exercise reduces cardiovascular disease risk, but how this protection is mediated at the molecular level remains incompletely understood.

**Methods:** Using multi-omic data from MoTrPAC, we integrated 13,790 selected molecular features across eight cardiac omic layers from 50 female and male rats undergoing 1-8 weeks of endurance training, as well as sedentary controls. Multi-omic temporal programs were defined using MOFA/MEFISTO and compared to physiological adaptation, human cardiac disease signatures, causal cardiomyopathy genes, and cross-tissue responses.

**Results:** Seven temporal programs of coordinated molecular responses were identified across cardiac omic layers. Transient metabolic responses showed sex-dependent timing and generally preceded sustained structural changes with limited correspondence to skeletal muscle, while complement proteins declined with training. Among these, a sustained program was associated with improvement of VO₂max and accounted for 38 of 50 significant overlaps with human cardiac disease signatures, frequently with opposing molecular responses, including reduced *Fn1* and increased *Klhl40* transcript and protein abundance by 8 weeks. To provide mechanistic context, we examined 112 genes causally linked to inherited cardiac disease, finding significant training-associated changes in 72 genes at the transcript level and 57 at the protein level. Notably, FLNC, BAG3, and PKP2 protein abundance increased, contrasting with reduced-dosage mechanisms in their respective cardiomyopathies. Conversely, disease-aligned and disease-opposing chaperone responses suggested coordinated protein quality-control mechanisms that may contribute to exercise-mediated cardioprotection.

**Conclusions:** Endurance training induces coordinated, time-dependent molecular responses across cardiac pathways implicated in human disease, including both disease-opposing changes in known causal genes and disease-concordant stress responses.

**Clinical implications:** *What is new?:*

- Endurance training induces distinct cardiac temporal programs, including a sustained multi-omic response associated with improved cardiorespiratory fitness, while transient metabolic responses generally precede progressive structural remodeling.
- Female and male hearts engaged broadly shared metabolic pathways with sex-specific timing of mitochondrial protein and post-translational responses.
- Sustained programs account for most overlaps with human cardiac disease signatures, frequently with opposing molecular responses; increased FLNC and BAG3 further suggest potential compensation for reduced-product cardiomyopathy mechanisms.

*What are the clinical implications?:*

- Endurance training induces molecular responses that significantly overlap with cardiovascular disease signatures. Both disease-opposing and disease-aligned regulation may help distinguish potentially protective adaptations from pathological remodeling.
- Training upregulates several causal cardiomyopathy genes implicated in haploinsufficiency, suggesting potential compensation for reduced gene dosage across distinct genetic disease contexts.
- Female and male hearts have distinct temporal responses in excitation-contraction coupling, metabolism, and cytoprotection, highlighting potential sex-specific mechanisms of cardiac adaptation.

## Introduction

Cardiovascular diseases remain the leading cause of mortality and a major cause of disability worldwide,^1^ with heart failure and cardiomyopathies contributing substantially to this global burden.^2^ These diseases converge on maladaptive myocardial remodeling involving altered cardiac structure and extracellular matrix, impaired excitation–contraction coupling, and disrupted mitochondrial energetics and substrate metabolism, ultimately compromising cardiac function.^3,4^ In contrast, regular physical activity and exercise training substantially reduce cardiovascular risk and promote cardiovascular health,^5^ while inducing adaptive remodeling across many of the same cardiac systems disrupted in disease.^6^ Moreover, sex-specific differences in structural, electrical, and functional cardiac adaptation to exercise have been described in humans, but the molecular and temporal organization underlying these differences is poorly defined.^7,8^ Most molecular studies have typically examined a limited number of molecular layers and training endpoints, constraining resolution of how these responses are coordinated over time into an adaptive cardiac state. The Molecular Transducers of Physical Activity Consortium (MoTrPAC) provides a unique framework to resolve these questions through longitudinal, sex-resolved multi-omic profiling of endurance training. Previous MoTrPAC analyses of multiple tissues and molecular layers from female and male rats trained for 1, 2, 4, or 8 weeks, with tissues collected 48 hours after an exercise bout, revealed widespread time- and sex-specific metabolic, mitochondrial, immune, and cellular-stress responses across the organism.^9^ Yet, the temporal architecture of cardiac molecular adaptation, and its relationship to physiological adaptation and human cardiac disease, has not been defined.

Here, we integrate eight molecular layers across the endurance-training time course in female and male rats to define the temporal organization of cardiac adaptation, resolve shared and sex-specific regulatory trajectories, and relate these exercise-associated molecular profiles to human cardiovascular disease signatures.

## Material & methods

### Study design and data sources

The MoTrPAC study included 60 six-month-old male and female Fischer 344 rats subjected to progressive treadmill endurance training for 1, 2, 4, or 8 weeks, with age-matched sedentary controls. Combined left and right ventricular tissue was collected 48 hours after the final exercise bout and rinsed in saline immediately after removal to preferentially capture molecular states maintained during training adaptation. Eight molecular layers were analyzed, with five to six animals per sex, time point, and assay. Assays included transcriptomics, DNA methylation, chromatin accessibility, global proteomics, lysine acetylation, phosphorylation, ubiquitination, and metabolomics. Experimental procedures, molecular profiling, and primary data processing followed published MoTrPAC protocols.^9^ Analyses used normalized molecular matrices and differential-analysis outputs (“rat-training-06” MoTrPAC collection c2.0), based on rat reference genome/proteome Rn7.2), including protein-corrected post-translational modification measurements. Physiological measurements, including maximal oxygen consumption (VO₂max), body composition, and circulating metabolic and hormonal markers, were obtained from MoTrPAC rat physiology datasets.^10^

### Single-omic pathway enrichment analysis

Sex- and time-specific responses were characterized using ranked Gene Ontology Biological Process enrichment with fgsea^11^ for nonmetabolomic layers, followed by clustering of related terms based on gene overlap and semantic similarity. Metabolites were analyzed separately by direction of change using KEGG-based network enrichment with FELLA.^12^

### Multi-omic factor analysis and temporal modeling

All eight molecular layers were integrated using MOFA+ (Multi-Omic Factor Analysis) with the MEFISTO (Method for Functional Integration of Spatial and Temporal Omics) temporal framework.^13,14^ Features showing nominal evidence of a training response (P≤0.01) were selected, with epigenetic features restricted to regulatory regions associated with responsive transcripts or proteins. The final dataset comprised 50 animals, including five per sex per experimental group. Female and male samples were modeled as separate groups with shared feature weights and sex-specific factor scores, using training duration as a continuous covariate. Seven factors were retained based on explained variance and cross-omic representation. Factor trajectories were standardized relative to sedentary controls and classified as sustained or transient. Factor-associated features were identified using MOFA weights and sex-specific correlations with factor scores, then characterized through Gene Ontology, KEGG, Reactome, and MitoCarta enrichment using assay-specific backgrounds. Enriched pathways were manually grouped into broader functional categories. Associations of features with physiological adaptations and cross-tissue conservation in skeletal muscle and liver were also evaluated.

### Human cardiovascular disease and pathogenic mechanisms

Cardiac transcriptomic and proteomic signatures from the Cardiovascular Disease Atlas^15^ were compared with exercise-associated temporal programs using rat–human ortholog mapping (see Supplementary Methods) and Fisher exact enrichment tests with assay-specific backgrounds. Overlapping responses were classified as disease-opposing or disease-aligned using the 8-week response for sustained programs and the mean 1–4-week response for transient programs. Cardiac gene–disease relationships and pathogenic mechanisms from Cardiac Gene2Phenotype,^16^ supplemented by ClinGen dosage-sensitivity evidence,^17^ were used to evaluate inherited cardiac disease genes. Exercise responses were classified as potentially compensatory, adverse, or mixed relative to established disease mechanisms.

### Statistical analysis

Analyses were stratified by sex unless otherwise indicated. Group comparisons used two-sided Wilcoxon rank-sum tests, and associations were assessed using Pearson or Spearman correlations, as appropriate. Nominal P≤0.01 was used to identify suggestive molecular responses for feature selection and exploratory analyses, whereas adjusted P≤0.05 generally defined statistical significance. Testing across multiple contrasts used the Benjamini–Hochberg procedure to control the false discovery rate (FDR ≤ 5%). Detailed analytical procedures, software versions, statistical thresholds, multiple-testing correction families, and analysis parameters are provided in the Supplementary Methods and figure legends. Analysis code is publicly available at GitHub (https://github.com/MoTrPAC/motrpac-rat-training-6mo-cardiac).

## Results

### Endurance training engages conserved cardiac molecular responses across sexes and omics layers

Cardiac profiles were obtained from 5–6 rats per sex and group (sedentary or 1-8 weeks of endurance training) across assays. Eight cardiac molecular layers were characterized, encompassing transcript abundance, epigenetic features (DNA methylation and chromatin accessibility), protein abundance, post-translational modifications (PTMs: acetylation, phosphorylation, and ubiquitination), and metabolites (**Figure 1A**). During the training period, cardiorespiratory fitness increased in both sexes, with VO₂max significantly elevated by 8 weeks (female FDR ≤ 1.4 × 10⁻⁶; male FDR ≤ 6.6 × 10⁻⁶; **Figure 1B**). As previously reported, endurance training was also accompanied by changes in body composition and circulating hormones, lipids and metabolites, with several responses differing between sexes (**Figure 1C**).^10,18^

**Figure 1.**
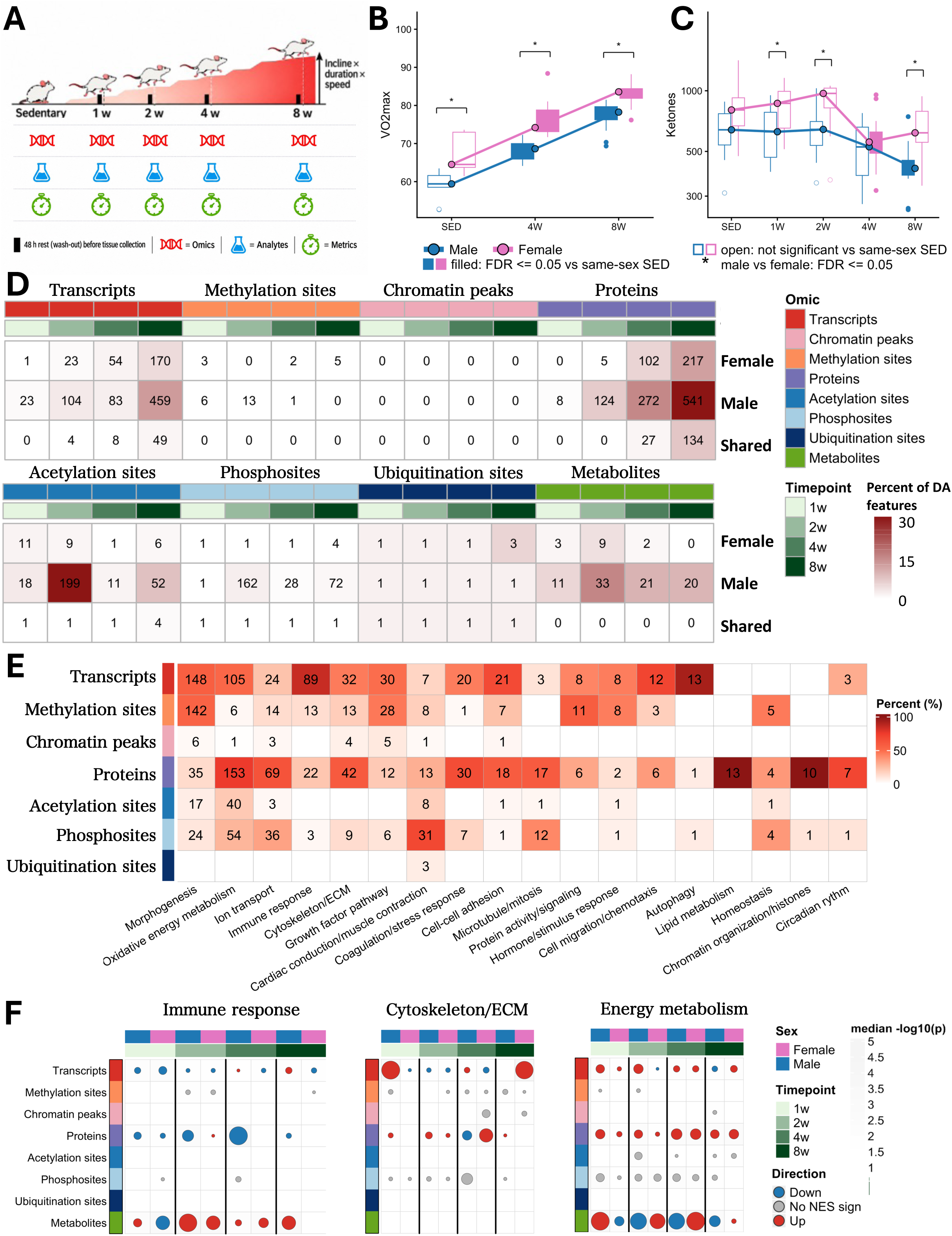
Endurance training induces heterogeneous molecular responses that converge on shared cardiac biological pathways. **A)** Overview of the endurance training study and cardiac multi-omic profiling. Male and female rats underwent progressive treadmill training for 1, 2, 4, or 8 weeks or remained sedentary (SED), with training workload progressively increased through incline, duration, and speed. Tissues were collected 48 hours after the final exercise bout. Cardiac profiling comprised transcriptomics, DNA methylation, chromatin accessibility, proteomics, acetylation, phosphorylation, ubiquitination, and metabolomics, together with physiological and circulating metabolic measurements. **B–C)** Physiological and circulating metabolic responses to endurance training. **(B)** VO₂max and **(C)** circulating ketone concentrations across the training course in females and males. Boxplots show individual animals, with sex-specific median trajectories overlaid. Filled boxes indicate significant differences from same-sex SED controls (FDR≤0.05); open boxes indicate nonsignificant differences. Asterisks indicate significant differences between females and males at the corresponding timepoint (FDR≤0.05). Comparisons were performed using two-sided Wilcoxon rank-sum tests with Benjamini-Hochberg correction. **D)** Number and proportion of significantly altered molecular features across omic layers, training timepoints, and sexes. Features were identified from timewise differential analyses comparing each training timepoint with SED controls at FDR≤0.05. Rows indicate female, male, and shared significant features between sexes at the same timepoint. Values indicate feature counts, and color intensity represents the percentage of quantified features within each molecular layer. **E)** Functional enrichment across molecular layers. Gene Ontology Biological Process enrichment was performed separately by omic layer, sex, and training timepoint using preranked GSEA, and significant terms (FDR≤0.05) were grouped into nonredundant biological categories based on gene-set overlap and semantic similarity. Categories containing ≥10 enriched terms are shown. Values indicate the number of significant terms within each category and molecular layer, and color intensity represents their percentage among terms assigned to that category. **F)** Temporal and sex-specific enrichment of selected biological programs across molecular layers. Columns represent female and male training timepoints and rows indicate molecular layers. Bubble size represents median enrichment significance. Red and blue indicate positive and negative enrichment, respectively; gray indicates enrichment without assigned direction. Epigenomic and post-translational modification layers were treated as direction-neutral.

Transcript and protein responses increased with training duration, whereas post-translational modifications (PTMs) and metabolite responses were prominent earlier (**Figure 1D**). Although males displayed a greater number of significant features, additional analyses revealed sex-concordant directional agreement between nominally responsive features across several molecular layers, particularly at later timepoints, despite differences in response magnitude (**Supplementary Results**, **Supplementary Figure 1**). Ranked gene-level enrichment identified shared biological responses involving oxidative energy metabolism, cardiac contraction, cytoskeletal and extracellular-matrix organization, lipid metabolism, immune regulation, and chromatin-associated processes (**Figure 1E**, **Supplementary Figure 2A**). Immune, cytoskeletal, and immune-related programs were among those represented across multiple molecular layers and in both sexes, although their magnitude, direction, and temporal dynamics varied (**Figure 1F**, **Supplementary Figure 2B**). Thus, training engaged shared cardiac pathways with sex-specific molecular and temporal regulation.

### Multi-omic integration resolves coordinated temporal programs of cardiac adaptation

To determine how these molecular responses were coordinated across omic layers and training duration, we integrated 13,790 features showing evidence of training responsiveness (nominal p-value ≤ 0.01) across eight molecular layers using MOFA+/MEFISTO. Seven factors were retained to capture the dominant cross-omic variation while limiting omic-specific components (**Supplementary Results**, **Supplementary Figure 3**). Factor trajectories and hierarchical clustering revealed a sustained response in Factor 2, characterized by a progressive increase throughout training in both sexes, whereas Factors 3 and 4 showed transient responses that peaked early before returning toward baseline (**Figure 2A–C, Supplementary Tables 1-2**). The remaining factors exhibited greater sex-specific differences in the timing, direction, or persistence of their responses. Factor 2 was most strongly and consistently associated with physiological adaptations, including increased VO₂max in both sexes and reductions in body mass and adiposity, particularly in males (**Figure 2D**). This program also showed the broadest representation across molecular layers, capturing transcriptomic, proteomic, post-translational, and metabolic variation (**Figure 2E**). At 8 weeks, fold changes of matched Factor 2-associated transcripts and proteins correlated in both sexes (r = 0.55, **Figure 2F**), highlighting conserved adaptation programs. Pathway enrichment linked the sustained Factor 2 program to cardiac structural and mechanical organization, as well as complement and coagulation pathways whose associated proteins decreased as Factor 2 increased (**Figure 2G**, **Supplementary results, Supplementary Table 3**). Notably, cytoskeletal pathways were enriched across transcriptomic, proteomic, and phosphoproteomic layers despite differences in regulatory direction, with concordant transcript–protein changes in *Klhl40* and *Ncam1* but discordant regulation of *Fhl1*.

**Figure 2.**
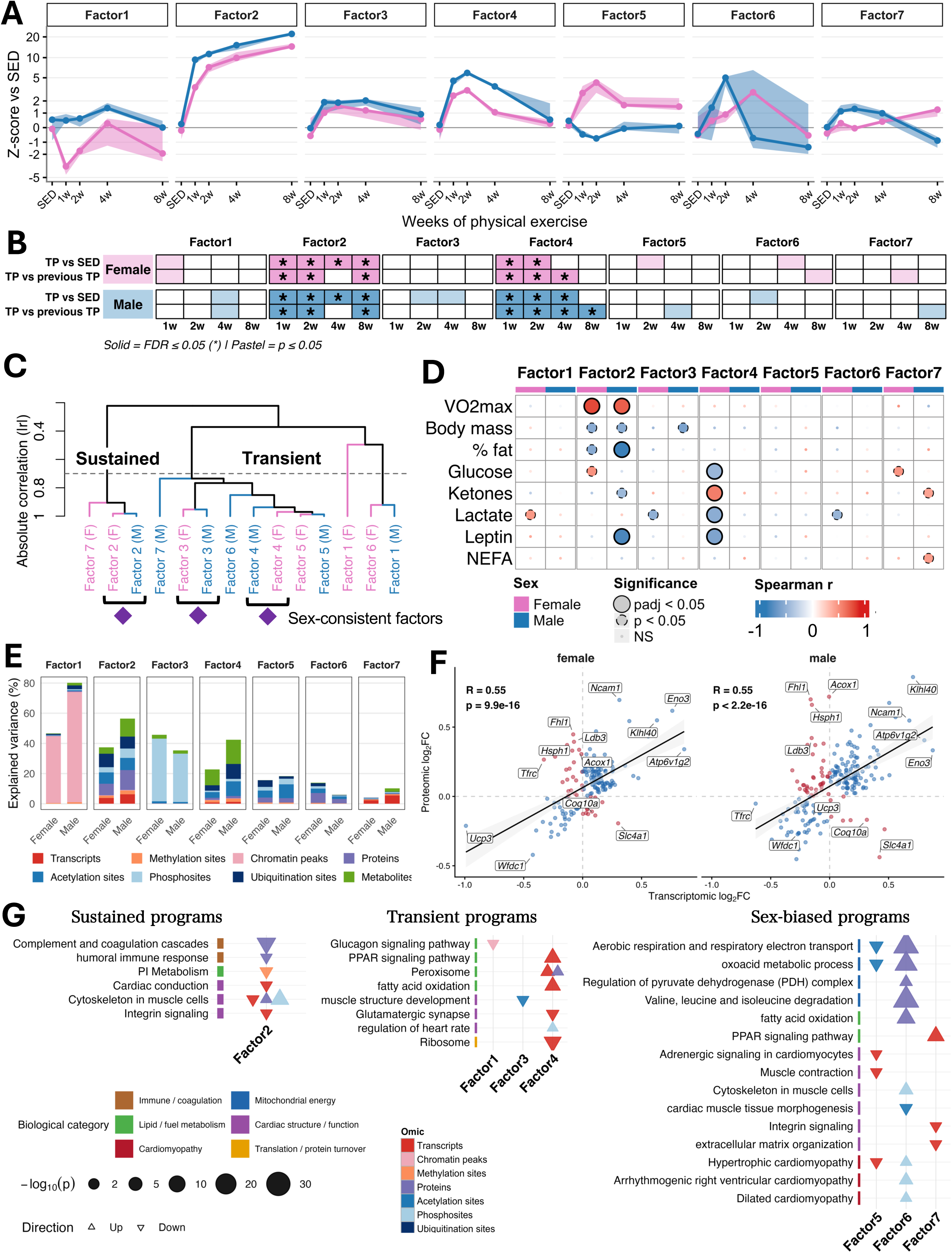
Multi-omic integration resolves seven temporal programs of cardiac adaptation. **A)** Temporal trajectories of the seven MOFA factors across the 8-week training course. MOFA sample scores were standardized within each sex and factor relative to sedentary (SED) animals and are shown as median z-scores with interquartile ranges (shading). Female and male trajectories are shown in pink and blue, respectively. Factor signs were oriented for biological interpretability. **B)** Significant changes in factor scores across the training course. For each factor and sex, exercise timepoints were compared with SED (top row) or with the preceding timepoint (bottom row) using Wilcoxon rank-sum tests. Pastel shading indicates nominal significance (P≤0.05); darker shading with an asterisk indicates significance after Benjamini-Hochberg correction (FDR≤0.05). **C)** Hierarchical clustering of sex-specific factor trajectories. Pearson correlations were calculated between median SED-normalized z-score trajectories, followed by average-linkage hierarchical clustering using 1−|r| as the distance metric. The dashed line indicates a correlation threshold of |r|=0.7. Branches are annotated as sustained or transient according to their temporal behavior, and purple diamonds indicate female and male trajectories from the same factor that cluster together above this threshold. **D)** Associations between factor and physiological and circulating metabolic traits. MOFA sample scores were correlated with physiological traits separately by factor and sex using Spearman correlation. Bubble color indicates Spearman ρ (blue, negative; red, positive), and bubble size and outline indicate statistical significance (large solid circles, FDR≤0.05; smaller dashed circles, nominal P≤0.05; small circles, nonsignificant). Female and male factor columns are indicated in pink and blue, respectively. VO₂max indicates maximal oxygen consumption; NEFA, nonesterified fatty acids. **E)** Stacked barplots showing the percentage of variance explained (R²) by each factor across molecular layers and sexes. Each segment represents variance explained within an individual molecular layer. **F)** Scatterplots comparing transcriptomic and proteomic log2(FC) at 8 weeks for Factor 2– associated genes. Genes were retained when significantly correlated with the Factor 2 trajectory in the corresponding sex (adjusted P≤0.05, |r|≥0.4) and quantified at both transcript and protein levels. Dot colors indicate concordant or discordant direction of regulation between molecular layers. Lines indicate linear regression fits; Pearson correlation coefficients and P values are shown. **G)** Enrichment summary of selected biological processes across factors, organized by sustained, transient, and sex-biased temporal programs. Enrichment was assessed among factor-associated molecular features using the corresponding quantified feature background. Columns indicate factors and rows indicate selected enriched pathways or biological processes. Dot color denotes molecular layer, shape indicates enrichment direction, and size represents −log10(P). Side annotations group terms into broad biological functions.

Sex-consistent transient programs involved early lipid utilization, PPAR signaling, ribosomal processes, and selected muscle-structural pathways (**Figure 2G**). In contrast, programs with sex-specific trajectories showed prominent enrichment of mitochondrial oxidative metabolism, paralleling sex-specific associations between factor trajectories and circulating metabolic markers (**Figure 2D**). Proteomic responses involving fatty-acid and branched-chain amino-acid catabolism, TCA-cycle processes, and respiratory electron transport were apparent at 2 weeks in males and from 4 weeks in females, whereas Factor 5 followed opposing trajectories between sexes, with prominent contributions from transcriptomic and acetylation-associated features. Taken together, these findings identify a sustained, multi-omic program of cardiac adaptation associated with improved fitness in both sexes, alongside transient and sex-specific responses reflecting distinct patterns of metabolic remodeling.

### Sustained cardiac programs frequently oppose human disease signatures and engage inherited cardiomyopathy genes

Given prominent cardiac structural and mechanical responses to training, we compared exercise-responsive programs with human cardiac transcriptomic and proteomic signatures from the Cardiovascular Disease (CVD) Atlas.^15^ These signatures encompassed 12 cardiovascular disease categories, including cardiomyopathies and heart-failure phenotypes, with broad rat orthologs coverage (**Supplementary Figure 5**). Exercise-associated programs significantly overlapped signatures from 11 of 12 disease categories, yielding 50 significant overlaps across sex-, ome-, and temporal-program-specific comparisons (**Figure 3A**). Sustained programs accounted for 76% of these overlaps, with particularly strong proteomic associations with dilated cardiomyopathy in both females (FDR = 3.15 × 10⁻¹¹) and males (FDR = 1.34 × 10⁻¹³), alongside significant overlaps with signatures of ischemic and hypertrophic cardiomyopathies and multiple heart-failure phenotypes (**Supplementary Table 4**). Significant phosphoproteomic overlaps included heart failure with reduced ejection fraction (HFrEF) in females and males (FDR = 9.18 × 10⁻⁴ and 2.30 × 10⁻³, respectively). Importantly, overlap did not imply concordant direction of effect: across sustained programs, 55% of shared transcripts and 62% of shared proteins, on average, were disease-opposing between exercise and disease. Patterns of exercise-disease discordance varied across diseases but were largely similar between sexes (**Figure 3A**).

**Figure 3.**
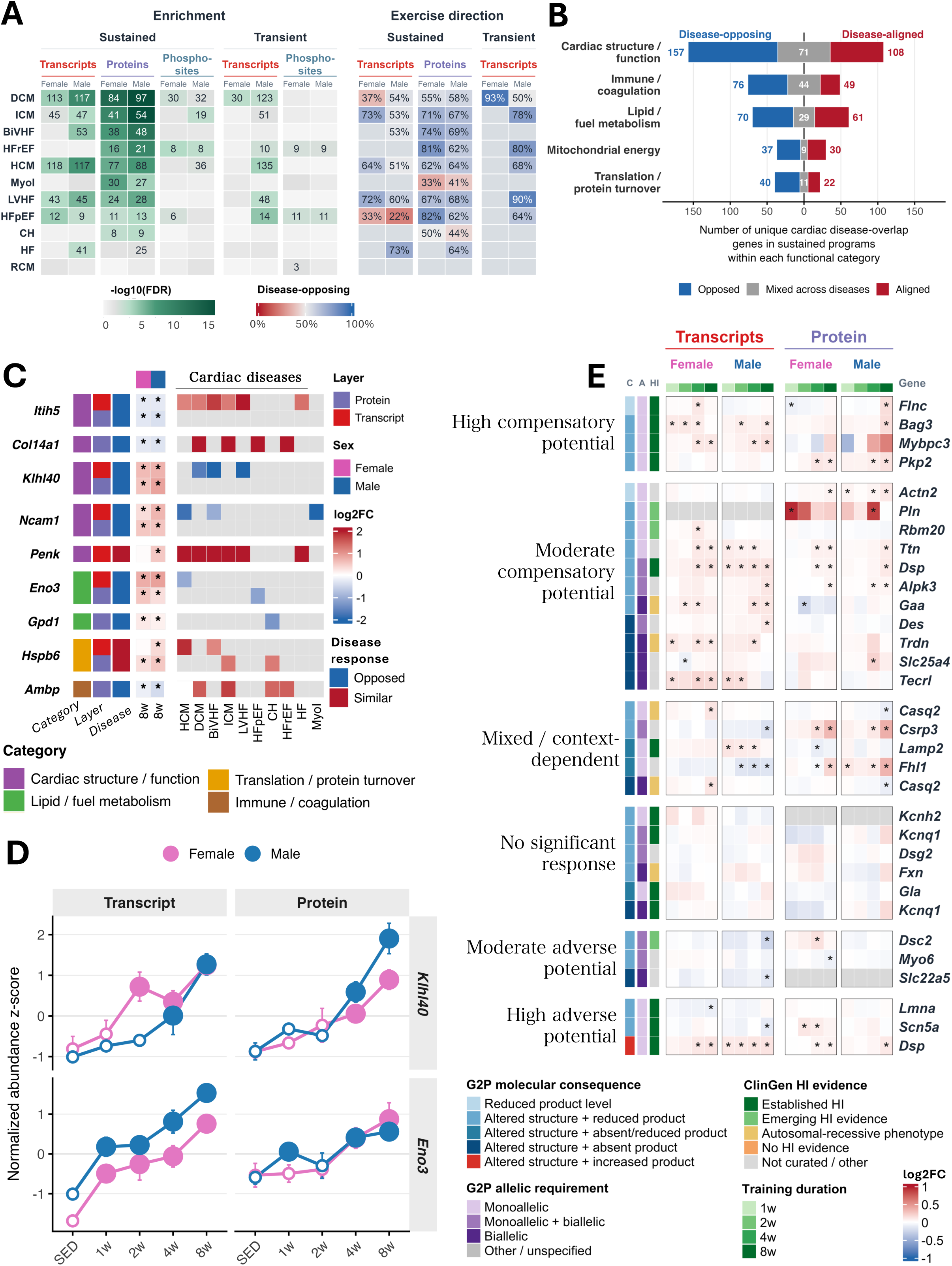
Sustained cardiac programs frequently oppose human disease signatures and involve genes causally linked to inherited cardiac disease. **A)** Combined heatmap showing overlap between exercise-responsive temporal programs and human cardiac disease signatures from the CVD Atlas. The left heatmap shows enrichment of factor-associated genes in disease-associated gene or protein sets, assessed by one-sided Fisher exact test and stratified by temporal program, molecular layer, and sex. Color indicates −log10(FDR), and values indicate the number of overlapping genes. The right heatmap shows, for transcriptomic and proteomic abundance layers, the percentage of overlapping genes regulated in opposite directions by exercise and cardiovascular disease. DCM: Dilated cardiomyopathy; ICM: Ischemic cardiomyopathy; BiVHF: Biventricular heart failure; HFrEF: Heart failure with reduced ejection fraction; HCM: Hypertrophic cardiomyopathy; Myol: Myocardial ischemia; LVHF: Left ventricle heart failure; HFpEF: Heart failure with preserved ejection fraction; CH: Compensated hypertrophy; HF: Heart failure; RCM: Restrictive cardiomyopathy. **B)** Diverging barplot summarizing gene-level counts of transcriptomic and proteomic features shared between sustained exercise programs and cardiovascular disease signatures. Features were collapsed by unique gene name, grouped into previously defined broad biological functions, and classified as disease-opposing, disease-aligned, or mixed according to the direction of their exercise response across disease contexts. **C)** Heatmap showing selected exercise-responsive genes. The left block reports transcript and protein log2(FC) at 8 weeks relative to sedentary controls in females and males; asterisks indicate p-value≤0.01. The right block presents median CVD Atlas log2(FC) across available datasets for the indicated cardiac disease categories. Row annotations indicate molecular layer, biological function, and exercise-disease response pattern. **D)** Temporal profiles of feature-wise z-scored transcript and protein abundance for Klhl40 and Eno3 across training timepoints and sexes. Lines and dots indicate mean±SEM across groups. Closed symbols indicate timepoints differing from sedentary controls at nominal p-value≤0.01. **E)** Heatmap representing exercise-induced regulation of 29 Cardiac Gene2Phenotype genes. Cells show RNA and protein log2(FC) relative to sedentary controls across training timepoints and sexes; asterisks indicate nominal p-value≤0.01. Relationships are grouped by their inferred potential relative to the curated disease mechanism: strong compensatory, compensatory, mixed or context-dependent, no significant response, adverse, or strong adverse. Annotations indicate G2P molecular consequence and allelic requirement and ClinGen haploinsufficiency evidence.

To identify biological processes underlying exercise-disease overlaps, we examined the functional organization of shared genes (**Figure 3B**). Cardiac structure/function comprised the largest class (157 disease-opposing, 108 aligned, 71 mixed directionality across diseases). Disease-opposing genes spanned extracellular matrix organization (*Itih5*, COL14A1), sarcomeric integrity (*Klhl40*), and cell adhesion and signaling (*Ncam1*). Disease-opposing responses also extended to lipid and fuel metabolism, mitochondrial energy metabolism, immune/coagulation pathways, and translation/protein turnover, with representative features including *Eno3*, GPD1, and AMBP (**Figure 3C-D**). Conversely, recurrent disease-aligned responses included *Hspb6* (cardiomyocyte stress protection) and *Penk* (neuropeptide signaling), suggesting that exercise and cardiac disease engage shared stress-responsive mechanisms.

Although these comparisons identified disease-opposing and disease-aligned molecular responses, their potential implications for inherited cardiac disease depend on the underlying pathogenic mechanisms. We therefore examined exercise responses across 112 genes causally linked to inherited cardiac diseases, curated from Cardiac Gene2Phenotype.^16^ Endurance training significantly altered 72 transcripts and 57 proteins at one or more timepoints (p-value≤0.01), with the number of regulated genes increasing throughout training to 51 transcripts and 38 proteins at 8 weeks. Using ClinGen dosage-sensitivity evidence^17^ and curated pathogenic mechanisms, we retained 32 gene–disease relationships involving 29 unique genes with reduced-product or increased-activity pathogenic mechanisms (**Figure 3E, Supplementary Table 5**). We then assessed whether exercise-induced changes might oppose (compensatory potential) or reinforce (adverse potential) these mechanisms. Potentially compensatory relationships outnumbered adverse ones (15 versus 6), or showed no significant response. These classifications likely reflect molecular effects rather than predicted clinical outcomes.

Four genes associated with haploinsufficient cardiomyopathies showed particularly notable, potentially compensatory exercise responses: *Flnc* (cytoskeletal integrity) and *Bag3* (proteostasis and myofibrillar stress adaptation) in dilated cardiomyopathy; *Mybpc3* (sarcomeric function) in hypertrophic cardiomyopathy; and *Pkp2* (desmosomal coupling) in arrhythmogenic right ventricular cardiomyopathy (**Figure 3E**). Their transcript and/or protein abundance increased by 4–8 weeks of training, with similar directional trends across sexes. *Dsp* illustrated a more context-dependent relationship: its induction at both transcript and protein levels might oppose reduced DSP abundance in some forms of arrhythmogenic cardiomyopathy but align with increased DSP abundance in dilated cardiomyopathy.

Together, these findings show that sustained cardiac adaptations to endurance training frequently opposed human cardiac-disease signatures. Further, exercise engaged causal cardiomyopathy genes in directions consistent with potential compensation for reduced gene dosage in specific disease contexts.

### Cardiac structural molecular changes increase with training while excitation– contraction regulation follows distinct trajectories

To characterize the largest disease-overlapping class, we mapped exercise-responsive features across an interconnected cardiac network spanning the extracellular-matrix (ECM), costamere/Z-disc, sarcomere, intercalated disc, and excitation–contraction machinery involved in force generation and transmission, cell–cell coupling, and electrical excitation (**Figure 4A–B**). Within this broader network, disease overlap was particularly concentrated in the ECM, where adhesion components were predominantly reduced with training. Disease-opposing responses included reductions in *Fn1*, *Sparc*, *Svep1*, and *Sned1*, alongside nine additional ECM features showing aligned or mixed relationships with human disease signatures. Intracellular structural responses involved predominantly increased abundance of costamere and Z-disc components (*Pdlim3*, *Csrp3*, *Nrap*, *Xirp2*, *Ldb3*) and sarcomeric proteins (*Myom2*, *Mybpc3*, *Ttn*, *Lmod2*). Changes in CTNNA1, CTNNB1, and *Dsp* implicated intercalated-disc adherens and desmosomal junctions, while increases in DMD indicated additional regulation of membrane–cytoskeletal coupling and sarcolemmal stability. Across structural compartments, 37 of 43 curated genes (86%) showed larger changes at 4–8 than at 1–2 weeks in at least one sex and omic layer, consistent with progressive regulation of cardiac structural components.

**Figure 4.**
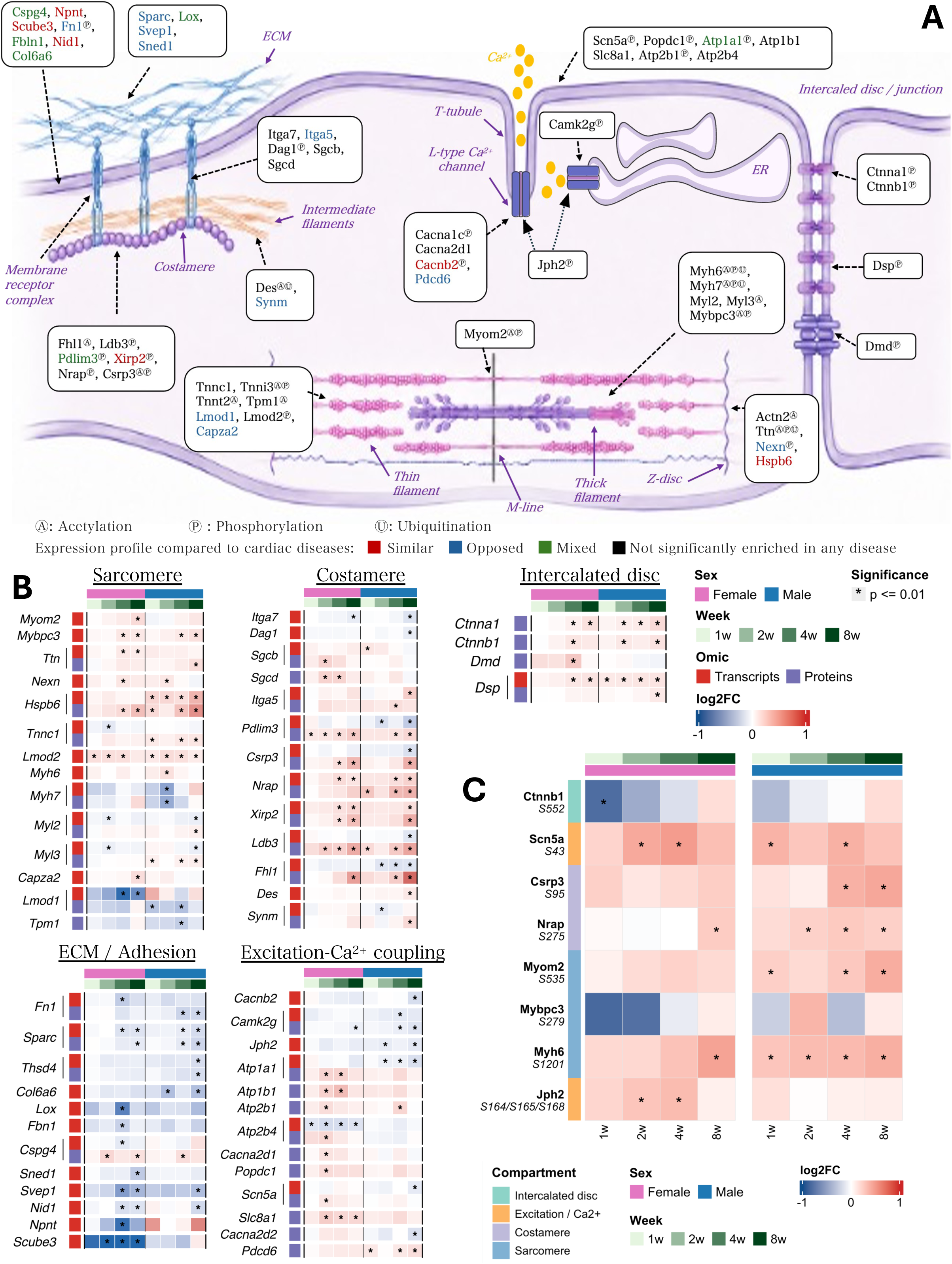
Endurance training coordinates structural, excitation–contraction, and post-translational remodeling across cardiomyocyte architecture. **A)** Schematic organization of exercise-responsive molecular features across cardiomyocyte compartments, including extracellular matrix (ECM) and adhesion, costamere/Z-disc, sarcomere, intercalated disc/cell junctions, and excitation–contraction and Ca²⁺-handling machinery. Boxes indicate genes or proteins exhibiting training-associated regulation, with superscripts denoting the molecular layer(s) in which regulation was detected (A: Acetylation, P: Phosphorylation, U: Ubiquitination). Text colors indicate the relationship between the exercise response and corresponding human cardiovascular disease signatures: opposed, similar, mixed, or not significantly enriched in the disease comparison. **B)** Temporal regulation of representative features from the functional compartments shown in A, displayed separately for females and males across 1, 2, 4, and 8 weeks of training relative to sedentary controls. Heatmap colors indicate log₂ fold change relative to sex-matched sedentary animals; asterisks indicate nominal p-value ≤ 0.01. Features are grouped by cardiomyocyte compartment. The annotation bars indicate sex, training duration, and molecular layer. **C)** Temporal regulation of selected phosphorylation sites spanning intercalated-disc, excitation– contraction, costamere/Z-disc, and sarcomeric proteins. Heatmap colors indicate log₂ fold change relative to sex-matched sedentary controls, and asterisks indicate nominal p-value ≤ 0.01. The left annotation denotes the cardiomyocyte compartment assigned to each phosphosite. Female and male responses are shown separately across 1, 2, 4, and 8 weeks of training.

A coordinated female-biased response emerged at 2–4 weeks that affected proteins involved in excitation–contraction and sarcolemmal ion-handling (**Figure 4B**). Affected proteins included Na⁺/K⁺-ATPase subunits (*Atp1a1*, ATP1B1), plasma-membrane Ca²⁺ pumps (ATP2B1, *Atp2b4*), the Na⁺/Ca²⁺ exchanger (SLC8A1), an L-type Ca²⁺ channel regulator (CACNA2D1), and proteins involved in membrane excitability and conduction (POPDC1, *Scn5a*). Notably, the transient increase in Scn5a protein abundance in females suggests a potentially compensatory molecular response for reduced gene dosage in haploinsufficiency-associated disease (**Figure 3E**). Male sarcolemmal protein responses were more heterogeneous and included decreases in junctional and excitation– contraction components such as *Cacnb2*, *Camk2g*, and *Jph2*. The temporal organization of this intracellular structural adaptation was largely cardiac-specific, as corresponding molecular features rarely followed similar trajectories in gastrocnemius skeletal muscle (Supplementary Results). PTM responses affected intracellular mechanical and electrical machinery, with phosphorylation predominating and acetylation particularly apparent among sarcomeric proteins (Figure 4A,C). Selected phosphosites mapped to proteins involved in force generation (MYH6, MYBPC3), Z-disc organization and mechanosensing (MYOM2, NRAP, CSRP3), membrane excitability and dyadic coupling (SCN5A, JPH2), and junctional signaling (CTNNB1). Together, these findings indicate that broadly sustained structural adaptation at the transcript and protein levels was accompanied by temporal and sex-specific PTM regulation.

### Cardiac metabolic responses exhibit sex-specific temporal and regulatory patterns

Compared with predominantly sustained structural responses, cardiac metabolic remodeling showed more diverse temporal and sex-specific patterns (**Figure 5A**, **Supplementary Figure 7**). Early lipid-handling responses involved fatty-acid delivery and processing (*Lpl*, FABP3, ACOX1), whereas APOC1–3 and APOE declined progressively, representing sustained, disease-opposing changes (**Supplementary Figure 8B**). Pyruvate dehydrogenase components (*Pdha1*, DLAT) showed a transient increase at 2 weeks in males, whereas the corresponding response in females peaked at 4 weeks. Oxidative proteins involved in β-oxidation (*Acadvl*), the TCA cycle (SDHA), and oxidative phosphorylation (*Ndufs1*), showed similar timing, with female responses emerging at 4 weeks. These responses generally attenuated by 8 weeks, unlike the more sustained oxidative gene and protein responses in skeletal muscle (**Supplementary Figure 7**). Because protein abundance alone does not capture post-translational regulation of metabolic enzymes, we examined their phosphorylation and acetylation (Figure 5B). These modifications were widespread among mitochondrial β-oxidation and TCA-cycle enzymes, with generally earlier and more pronounced responses in males, suggesting sex-dependent post-translational regulation of cardiac metabolism. In females, PDHA1 showed early increases in S293 phosphorylation and later K244 acetylation, whereas males showed early Y242 phosphorylation. Male responses also included increased HADHA K728 acetylation at 2–4 weeks, alongside multiple modifications of TCA-cycle enzymes.

**Figure 5.**
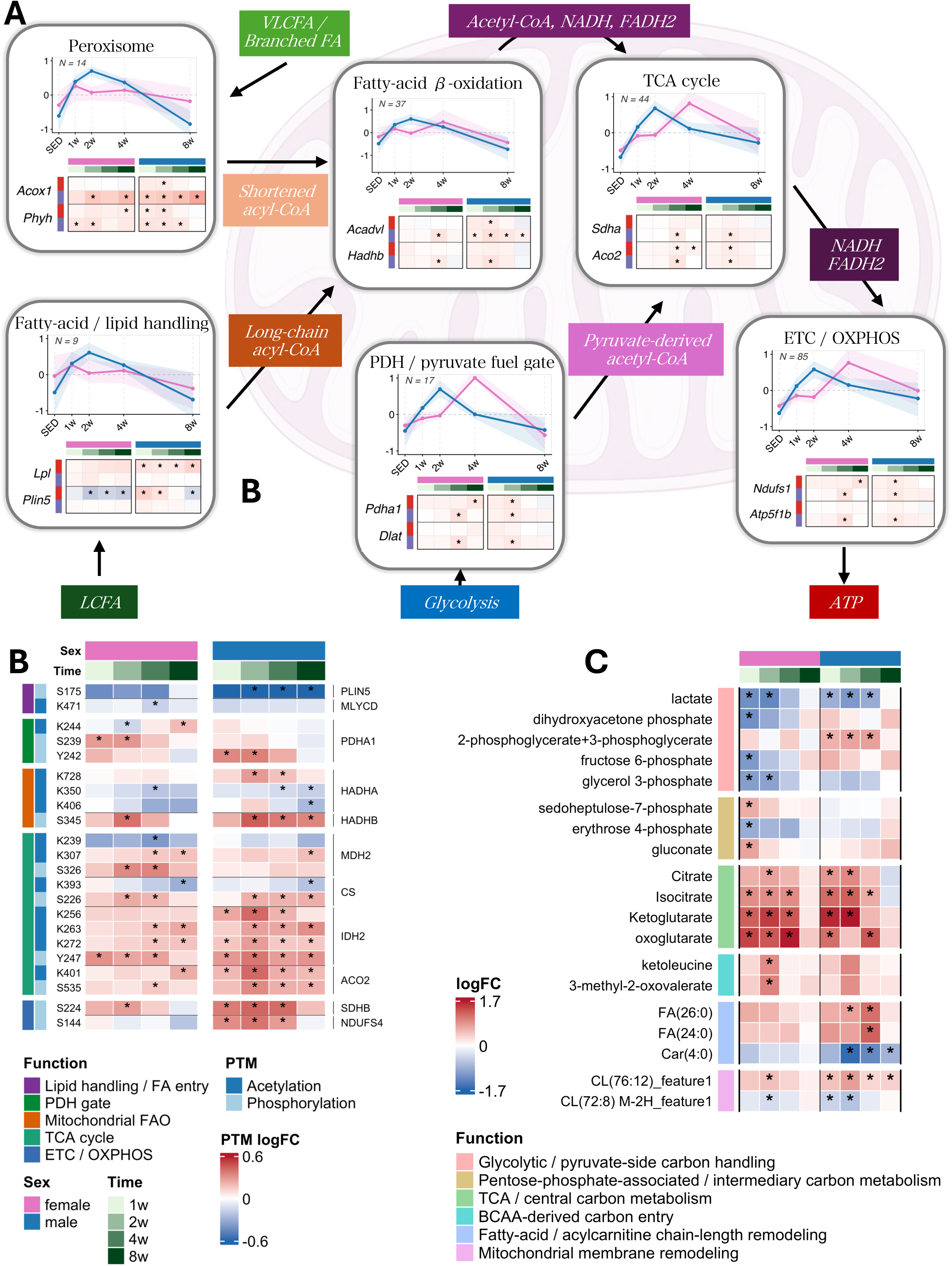
Multi-layer remodeling of cardiac fuel metabolism during endurance training. **A)** Temporal organization of transcript and protein abundance across major fuel-handling and mitochondrial pathways in heart and gastrocnemius skeletal muscle (SKMGN). Pathways are arranged according to the flow of substrates through fatty-acid/lipid handling and peroxisomal metabolism, mitochondrial fatty-acid β-oxidation, pyruvate dehydrogenase (PDH)–mediated pyruvate entry, the tricarboxylic acid (TCA) cycle, and electron transport/oxidative phosphorylation (ETC/OXPHOS). For each pathway, normalized transcript and protein abundances were standardized as z-scores within tissue and sex across sedentary (SED) and all training timepoints. Lines show the mean trajectory across genes after averaging each feature within timepoint; shaded ribbons indicate the 25th–75th percentile across feature-level trajectories. Pink and blue indicate females and males, respectively. N indicates the number of distinct genes represented in each pathway. **B-C)** Temporal remodeling of selected protein-normalized post-translational modifications (**B**) and cardiac metabolites (**C**). Heatmap values indicate log2 fold change (log2FC) relative to sex-matched sedentary controls at each training timepoint. Columns represent 1, 2, 4, and 8 weeks of training separately for females and males; asterisks indicate nominal p-value≤0.01 versus sedentary controls. Functional, sex, training-duration, and molecular-layer annotations are indicated by the accompanying color bars.

Despite sex differences in mitochondrial protein abundance and PTM responses, metabolite changes occurred in the early training adaptation in both sexes (**Figure 5C**). At 1 week, lactate decreased, whereas citrate, isocitrate, and 2-oxoglutarate increased. Females showed reduced dihydroxyacetone phosphate and glycerol-3-phosphate and changes of pentose-phosphate intermediates, whereas males showed increases in phosphoglycerate intermediates. Cardiolipin species also showed reciprocal early changes, consistent with shifts in the mitochondrial lipid profile. Cardiac triglycerides also followed distinct trajectories, reaching their greatest reduction at 4 weeks in females but continuing to decline through 8 weeks in males (**Supplementary Figure 8C**). Together, these findings indicate that both sexes engage broadly overlapping cardiac metabolic pathways through distinct temporal and regulatory trajectories. In females, early PTM and metabolite changes preceded broader mitochondrial protein induction at 4 weeks, indicating that metabolic remodeling was already underway before these protein-abundance changes emerged.

### Training increases chaperone and redox proteins while selectively reducing complement–hemostatic components

Following the metabolic changes described above, we next examined whether training also engaged systems supporting protein quality control and redox homeostasis. Molecular chaperones (HSPA1B, HSPB1, HSPB6, HSPD1, HSPA9) increased progressively in heart, with similar responses in skeletal muscle and liver as previously reported (**Supplementary Figure 9**).^9^ Several of these are also elevated in cardiovascular disease, suggesting shared stress-associated molecular responses rather than disease-opposing changes. Mitochondrial redox-defense proteins (PRDX3, SOD2, MSRB2) showed earlier responses in males and more prominent induction beginning at 4 weeks in females, paralleling the timing of mitochondrial oxidative protein changes. A distinct female cytoprotective response involved CRYAB, GSTA4, and GLRX3 (**Supplementary Figure 9**), whereas immune-related transcriptional responses were more heterogeneous and showed limited cross-tissue concordance.

In contrast, extracellular complement and hemostatic proteins showed coordinated reductions, particularly in males. These included complement components and regulators (C3, CFH, MASP2) and proteins involved in contact activation, coagulation, and fibrinolysis (KLKB1, F12, PLG, CPB2), many of which opposed cardiac-disease signatures. These coordinated reductions were not consistently reproduced in gastrocnemius skeletal muscle or liver at corresponding training time points (**Supplementary Figure 9C**). The kallikrein–kinin system showed additional complexity, with KLKB1 and F12 decreasing while KNG1 and KNG2 increased later in training across heart, skeletal muscle, and liver, indicating differential regulation rather than uniform suppression of this pathway. Together, these findings reveal a layered cardiac stress-adaptation response, combining broadly shared stress-response programs also engaged in cardiac disease with sex- and tissue-dependent regulation of redox, immune, and hemostatic pathways.

## Discussion

The present study from MoTrPAC represents one of the most comprehensive analyses of the cardiac molecular response to exercise available. Despite prior evidence that endurance training engages molecular pathways implicated in pathological cardiac remodeling,^6^ these data clarify how these responses are coordinated across molecular layers and time. Here, time-course integration of eight cardiac molecular layers revealed a progressive adaptation organized across energetic, structural, excitation–contraction, and stress-response programs, with sustained responses preferentially overlapping human cardiomyopathy and heart-failure signatures. Thus, exercise regulates disease-relevant pathways with distinct timing, context, and often direction.

The clearest expression of this adaptive state was a progressive, sustained program shared by both sexes, showing the strongest association with increasing cardiorespiratory fitness and transcript–protein concordance at 8 weeks (r=0.55 in both sexes). It encompassed energetic, structural, electrical, and stress-response processes, including *Klhl40*, which stabilizes sarcomeric thin-filament proteins and thereby links the sustained response to maintenance of contractile integrity.^19^ *Eno3* provided a complementary metabolic example, with concordant transcript–protein induction across sexes and previously reported persistent cardiac elevation after endurance training in association with a cardioprotective phenotype.^20^ At the pathway level, this sustained state also included coordinated regulation of complement–contact and hemostatic systems, with reductions in C3, MASP2, F12, and KLKB1 that were particularly prominent in males, frequently opposed cardiac-disease signatures, and were not consistently reproduced in skeletal muscle and liver. Although hearts were rinsed in saline at collection, contributions from residual circulating proteins cannot be excluded, limiting the attribution of these changes exclusively to local cardiac regulation.

More broadly, sustained programs accounted for 38 of 50 significant overlaps with human cardiomyopathy and heart failure signatures. Across these programs, disease-opposing responses represented a modest majority of shared molecular signals: on average, 55% of transcriptomic and 62% of proteomic signals were regulated in opposite directions by exercise and disease, with similar proportions in females and males. Cardiac structure and function represented the largest disease-overlapping class, with responses spanning an interconnected network of ECM, costamere, Z-disc, sarcomeric, intercalated-disc, and excitation–contraction components involved in mechanical load sensing, force transmission, and contractile function.^21^ Although disease-opposing responses may highlight mechanisms counteracting pathological remodeling, disease-aligned responses may also reflect adaptive or compensatory processes, underscoring the context dependence of shared cardiac molecular programs.^22^ Both patterns may therefore offer potential therapeutic insights.

Within this network, training predominantly reduced ECM- and adhesion-associated components involved in matrix assembly and maturation, including *Fn1*, *Sparc*, *Lox*, and *Svep1*. Several of these reductions opposed pathological matrix remodeling: SPARC-dependent collagen maturation and LOX-associated cross-linking have both been implicated in myocardial stiffening.^23,24^ *Fn1* further illustrates the context dependence of this response, as its deletion attenuates pressure-overload hypertrophy yet does not impair exercise-induced physiological cardiac growth.^25^ These findings suggest selective regulation of matrix assembly and maturation rather than reversal of a uniform fibrotic program. The partial conservation of these responses in gastrocnemius, particularly in males, suggests that ECM adaptation extends across mechanically loaded striated tissues.^26^

In contrast to the partially conserved extracellular changes, components of the intracellular load-bearing network, including *Csrp3*, *Fhl1*, *Ttn*, *Nrap*, *Myom2*, and *Xirp2*, were predominantly upregulated with training, with responses largely specific to the heart. This network participates in cardiac mechanosensing and has been implicated in pathological remodeling, notably through the MLP–telethonin–titin complex and FHL1-associated titin signaling.^27–29^ Experimental models further show its involvement in both physiological and pathological adaptation: *Fhl1* deficiency attenuates pressure-overload hypertrophy, whereas titin N2B disruption blunts physiological hypertrophy induced by endurance exercise.^30^ A direct example of disease-aligned regulation emerged at the phosphorylation level: MYOM2 S535 and NRAP S275 phosphorylation increased progressively or remained elevated with training, particularly in males, and these same sites are hyperphosphorylated in experimental HFpEF.^31^ Together, these results suggest that exercise and pathological cardiac loading may engage overlapping mechanosensory and phosphoregulatory pathways, without necessarily producing similar functional consequences.

Beyond broad exercise–disease overlap, the regulation of causal cardiomyopathy genes offers more specific mechanistic insight. Using Cardiac Gene2Phenotype,^16^ we identified gene– disease relationships with more compensatory than adverse potential. Among those involving reported haploinsufficiency,^17^ *Flnc* and *Bag3* increased with training in both sexes, consistent with potential compensation for reduced gene dosage in dilated cardiomyopathy. BAG3 facilitates FLNC turnover at the sarcomere through HSP70 and small heat-shock proteins.^32^ Experimental restoration of BAG3 improves sarcomeric protein turnover and contractile function in failing hearts.^33^ Notably, HSPA1B and HSPB6, members of these respective chaperone families, increased with both training and cardiac disease, representing disease-aligned responses consistent with shared cytoprotective stress signaling.^34^ In contrast, HSPH1, an HSP70-associated co-chaperone involved in protein refolding and disaggregation, was induced by exercise but showed opposing regulation in cardiac disease.^35^ Together, these findings suggest that endurance training engages a coordinated protein quality-control response involving causal cardiomyopathy genes and broader chaperone regulation that may contribute to the cardioprotective effects of exercise.

The clinical implications of these potentially compensatory responses remain uncertain. A small study of FLNC truncating-variant carriers found no significant association between higher-intensity exercise and adverse cardiac outcomes.^36^ *Mybpc3* transcript abundance also increased with training, contrasting with the reduced cMyBP-C dosage implicated in hypertrophic cardiomyopathy. Experimentally, cMyBP-C deficiency exacerbates cardiac dysfunction under pressure overload.^37^ However, in carriers of truncating MYBPC3 variants, high exposure to dynamic exercise was associated with increased risk of malignant ventricular arrhythmias.^38^ Similarly, PKP2 protein abundance increased with training in both sexes, potentially opposing the reduced-product mechanism implicated in arrhythmogenic cardiomyopathy. PKP2 links desmosomal integrity to electrical excitability, with PKP2 loss reducing NaV1.5 localization and sodium current.^39^ Yet endurance training promotes Ca²⁺-handling instability and arrhythmias in PKP2-deficient mice, while greater endurance-exercise exposure in human desmosomal mutation carriers is associated with increased disease penetrance, ventricular arrhythmias, and heart failure.^40,41^ These clinical and experimental observations underscore that exercise-induced molecular changes consistent with opposing reduced gene dosage do not necessarily predict protection in genetically susceptible hearts. Whether such responses confer functional benefit will require direct testing in appropriate models.

Joint modeling identified shared programs with sex-specific trajectories, while individual proteins highlighted timing differences in excitation–contraction and metabolism. These findings complement previous reports of sex differences in human cardiac adaptation to exercise.^7,8^ Females showed prominent ion-handling changes at 2–4 weeks, followed by broader induction of mitochondrial oxidative proteins from 4 weeks onward. These processes are closely connected, as Ca²⁺ handling couples excitation–contraction to mitochondrial metabolism, a relationship disrupted in heart failure.^42^ Females also showed progressive increases in cytoprotective proteins, including GLRX3, whose deficiency disrupts both redox homeostasis and sarcoplasmic-reticulum Ca²⁺ handling before cardiac dysfunction develops.^43^ In males, oxidative and mitochondrial redox-defense proteins increased earlier, consistent with findings from an earlier analysis of the same rat cohort.^44^ However, this earlier oxidative induction coincided with increased HADHA K728 acetylation at 2 weeks, a modification shown to inhibit fatty-acid oxidation in experimental HFpEF.^45^ Cardiac triglycerides followed a different temporal pattern, reaching their greatest reduction at 4 weeks in females but continuing to decline through 8 weeks in males, alongside broader lipid changes in liver and adipose tissue.^18,46^

Despite sex-dependent mitochondrial protein trajectories, citrate, isocitrate, and 2-oxoglutarate increased transiently in both sexes. Levels of multiple cardiac oxidative proteins peaked during early or intermediate training before attenuating, whereas structural changes became more pronounced later. In addition, the pentose phosphate pathway, linked to exercise-induced NADPH production, physiological cardiac growth, and ischemic protection,^47^ showed early changes in associated metabolites in females. This broader temporal organization parallels studies in mice showing that changes in cardiac energy charge and glucose-derived TCA metabolism preceded physiological growth and subsequently attenuated,^48^ while experimental modulation of cardiac glycolysis influenced physiological growth.^49^ In contrast, oxidative gene and protein abundance in gastrocnemius increased more progressively and remained elevated throughout training.^50^ Together, these findings suggest that early metabolic responses may precede more sustained structural remodeling in the already highly oxidative heart.^51^ Whether these programs are conserved in human myocardium remains unknown.

Several limitations warrant consideration. Different animals were sampled at each training duration, so trajectories reflect group-level patterns and cannot establish causal sequence. Bulk ventricular profiling cannot distinguish cell-intrinsic regulation. Further, tissue collection 48 hours after the final bout favors training-associated responses but cannot exclude residual acute effects. Cardiac mass, chamber dimensions, and ventricular function were not assessed, so structural molecular changes cannot be linked directly to anatomical and functional adaptation. Similarly, VO₂max reflects integrated central and peripheral adaptations rather than cardiac function alone. Abundance and PTM levels do not establish protein activity, pathway flux, or functional effects. Rat-human differences in ventricular myosin composition^52^ and Ca²⁺ removal^53^ further constrain translation. Overlap with heterogeneous human disease signatures does not demonstrate protection, and responses in wild-type animals cannot establish compensation in pathogenic-variant carriers.

In summary, endurance training establishes a temporally organized cardiac adaptive state in which a sustained molecular program is associated with improvements in cardiorespiratory fitness and accounts for much of the overlap with human cardiac disease, often in the opposite direction to disease. Female and male hearts converge on broadly shared adaptations through differently ordered regulatory trajectories, while structural and excitation–contraction responses show less correspondence with skeletal muscle than metabolic adaptation. These findings highlight timing, direction, and regulatory context as key features distinguishing adaptive cardiac stress from pathological remodeling.

## Data and Code Availability

### Data availability

The multi-omic and phenotypic data analyzed in this study are publicly available through the MoTrPAC Data Hub (https://motrpac-data.org/).

### Code availability

Analysis code supporting the findings of this study is publicly available on GitHub (https://github.com/MoTrPAC/motrpac-rat-training-6mo-cardiac).

## Supplementary materials

### Supplementary methods

#### Study design, molecular profiling, and data processing

The original MoTrPAC cohort comprised 60 six-month-old male and female Fischer 344 rats subjected to progressive treadmill endurance training for 1, 2, 4, or 8 weeks, with age-matched sedentary animals serving as controls. At each training duration, combined left and right ventricular tissue was collected 48 h after the final exercise bout to reduce the contribution of acute post-exercise signaling and preferentially capture molecular states maintained during training adaptation. Eight molecular layers were profiled: transcriptomics by RNA sequencing, DNA methylation by reduced-representation bisulfite sequencing (RRBS), chromatin accessibility by assay for transposase-accessible chromatin using sequencing (ATAC-seq), mass-spectrometry-based global proteomics, lysine acetylation, phosphorylation, ubiquitination, and metabolomics, including lipid species. The nominal experimental design included five animals per sex and time point for transcriptomic, epigenetic, and metabolomic profiling and six animals per sex and time point for proteomic and post-translational modification (PTM) profiling.

Experimental procedures, endurance-training intervention, tissue collection, molecular profiling, quality control, normalization, and primary assay processing were performed according to standardized MoTrPAC protocols described previously.^9^ In the present study, all genomic and proteomic datasets were re-analyzed against the Rn7.2 rat reference genome and proteome. Proteomics data were searched using the cloud-based MS-GF+ proteomics pipeline. For PTM analyses, protein-corrected acetylation, phosphorylation, and ubiquitination measurements were used to account for variation in the abundance of the corresponding proteins before PTM-specific downstream analyses.

#### Feature annotation and cross-species mapping

Rat gene identifiers and human orthologs were assigned using org.Rn.eg.db (v3.22.0), biomaRt (v2.66.1), and the Rat Genome Database.^54^ Because some rat genes mapped to more than one potential human ortholog, mappings were not assumed to represent strictly one-to-one ortholog relationships. For epigenetic analyses, ATAC-seq and RRBS genomic regions were annotated using GenomicRanges (v1.62.1) and ChIPseeker (v1.46.1) against the rn7 RefSeq transcript annotation. Regions were classified as promoter, exon, intron, or intergenic, and distance to the transcription start site was retained for subsequent feature-selection analyses. For PTM analyses requiring human site annotation, rat protein sequences from the mRatBN7.2 reference proteome were aligned to the reviewed human UniProt proteome using BLASTp. Alignments with ≥60% sequence identity were retained, and the best human match was selected based on identical-residue and positive-match counts. Rat modification sites were projected through the protein alignment while accounting for sequence gaps, and mapped sites were retained only when the corresponding human residue was present and matched the expected amino-acid identity.

### Physiological measurements and statistical analysis

Physiological measurements were obtained from the MoTrPAC six-month rat physiology resources, MotrpacRatTraining6moData (v2.0.0) and MotrpacRatTrainingPhysiologyData (v2.0.0).^10^ Measurements included maximal oxygen consumption (VO₂max), body mass, percentage body fat, circulating glucose, ketones, lactate, nonesterified fatty acids, insulin, leptin, and corticosterone.

Post-intervention measurements were compared between each trained group and the corresponding sex-matched sedentary group using two-sided unpaired Wilcoxon rank-sum tests. Sex differences within each experimental group were assessed using the same test. Benjamini– Hochberg correction was applied separately to female-versus-sedentary, male-versus-sedentary, and female-versus-male comparison families across physiological variables and time points.

### Single-omic pathway enrichment analysis

#### Gene Ontology enrichment

For each nonmetabolomic molecular layer, sex, and training time point, ranked gene-set enrichment analysis was performed against Gene Ontology Biological Process terms obtained for Rattus norvegicus using msigdbr (v25.1.1). Features were ranked as sign(effect) × −log10(P). When multiple molecular features mapped to the same gene name, the feature with the largest absolute ranking statistic was retained. Gene-set enrichment was performed using fgsea (v1.36.2), restricting gene sets to 10–500 genes. Terms with adjusted P≤0.05 were considered significantly enriched.

To reduce redundancy among enriched GO terms, related terms were grouped using a combined measure of leading-edge gene overlap and ontology-based semantic similarity. Gene overlap was quantified using the overlap coefficient, and semantic similarity was calculated using the information-content-based Rel metric implemented in GOSemSim (v2.36.0) with org.Rn.eg.db (v3.22.0). Average-linkage hierarchical clustering was applied to the resulting similarity matrix, and clusters were defined using a similarity threshold of 0.10. Ubiquitination was excluded from the principal multi-omic GO clustering because of the limited number of responsive features. Cluster labels were manually assigned based on pathway content and representative terms.

#### Metabolite pathway enrichment

Metabolites with nominal P ≤0.01 were analyzed separately by sex, training time point, and direction of change. Metabolite annotations were mapped to KEGG compound identifiers using KEGGREST (v1.50.0), and network-based enrichment was performed using FELLA (v1.30.0) with a Rattus norvegicus KEGG network and the diffusion method with normality approximation. Pathways or network components with FELLA P-score ≤0.05 were retained. Related enriched pathways were grouped based on overlap among contributing metabolites and manually summarized into broader metabolic categories.

### Multi-omic factor analysis and temporal modeling

#### Feature selection for multi-omic integration

All eight molecular layers were included in the integrated analysis. Candidate features were defined as the union of features with nominal P ≤0.01 in at least one sex-specific timewise contrast or in the sex-specific overall-training test. To improve interpretability of the epigenetic layers, candidate features were additionally restricted to regions associated with genes exhibiting an RNA or protein training response. RRBS loci were required to fall within −1,000 to +1,000 bp of the transcription start site, whereas ATAC-seq regions were restricted to −1,500 to +1,000 bp. Features containing missing normalized values within their original molecular matrix were removed before cross-view alignment.

#### Multi-omic factor modeling

Samples represented in at least seven of the eight molecular layers were retained, resulting in a final integrated dataset of 50 animals, with five animals per sex in each of the sedentary, 1-, 2-, 4-, and 8-week groups. Multi-omic integration was performed using MOFA+ (v1.20.2)^13^ with the MEFISTO^14^ continuous-covariate framework. Female and male samples were modeled as separate groups with shared feature weights and sex-specific factor scores, and training duration was included as the continuous temporal covariate. Molecular views were scaled before model fitting. To determine the latent dimensionality, models containing 2–15 factors were fitted using 10 independent random seeds for each factor number. Factors were considered to capture substantial molecular variation when they explained ≥2% of variance in at least two molecular views and ≥5% cumulatively across views, or ≥5% in any individual molecular view. A final model of 7 factors was selected to balance cross-omic variance explained with the number of factors dominated by a single molecular view.

#### Factor trajectories and temporal classification

Factor scores were extracted using MOFA+ and standardized within each sex relative to the mean and standard deviation of sedentary animals. Temporal trajectories were summarized by the median and interquartile range at sedentary, 1-, 2-, 4-, and 8-week time points. Differences among experimental groups were assessed using two-sided unpaired Wilcoxon tests, with Benjamini– Hochberg correction across pairwise comparisons within each factor and sex.

Similarity among temporal factor trajectories was assessed using Pearson correlations between sex-specific median factor scores across the five experimental time points, and factors were grouped according to their temporal dynamics into sustained and transient response patterns.

#### Associations of multi-omic factors and physiological adaptation

Associations between multi-omic factor scores and physiological variables were assessed separately by sex using Spearman rank correlations. VO₂max, body mass, and percentage body fat were expressed as post-intervention minus pre-intervention changes, whereas circulating analytes were represented by post-intervention measurements. P values were adjusted across physiological variables within each factor and sex using the Benjamini–Hochberg procedure.

#### Identification of factor-associated molecular features

For each factor and molecular view, features were ranked by absolute MOFA weight, and the smallest set accounting for 50% of the cumulative absolute weight was retained. Because feature weights were shared across sexes whereas factor contributions could differ between females and males, candidate features were subsequently correlated with individual factor scores separately by sex using Pearson correlation to identify sex-specific feature–factor associations. P values were adjusted using the Benjamini–Hochberg procedure within each factor, molecular view, and sex, and features with |r| ≥0.4 and adjusted P ≤0.05 were considered significantly associated with the corresponding factor. Factor-associated features were further separated according to the sign of their MOFA weight. Weight direction reflects association with the latent factor trajectory and was therefore not interpreted directly as increased or decreased abundance with training.

#### Pathway enrichment of factor-associated features

To broadly characterize the biological processes captured by each factor, factor-associated features were pooled across sexes and analyzed separately for each factor and molecular layer. Features were further separated according to the sign of their MOFA weight. Overrepresentation analyses were performed on unique rat genes using the genes measured in the corresponding molecular layer as the assay-specific background. KEGG, Reactome, and Gene Ontology Biological Process enrichment was performed using gprofiler2 (v0.2.4) with *Rattus norvegicus* as the target organism. Mitochondrial pathway enrichment was additionally performed using MitoCarta 3.0 gene sets with clusterProfiler (v4.18.4). Enriched pathways were manually grouped into broader biological categories for visualization and interpretation.

#### Human cardiovascular disease enrichment

Human cardiovascular disease signatures were obtained from the Cardiovascular Disease (CVD) Atlas^15^ and restricted to transcriptomic and proteomic datasets derived from cardiac tissues. Disease contexts included cardiomyopathies, ischemic heart disease, cardiac hypertrophy, and heart failure phenotypes. Human gene and protein identifiers were harmonized and mapped to rat molecular features using the rat-to-human ortholog annotations described above. For each sex, molecular layer, and temporal-response class, enrichment of factor-associated genes in individual human cardiovascular disease signatures was assessed using one-sided Fisher exact tests. Human orthologs quantified in the corresponding rat molecular layer were used as the assay-specific background, and P values were adjusted within each molecular layer using the Benjamini– Hochberg procedure. Exercise responses overlapping human cardiovascular disease signatures were classified as disease-opposing or disease-aligned according to the direction of change, using the 8-week log-fold change for sustained responses and the mean log-fold change across 1, 2, and 4 weeks for transient responses. Directional comparisons were restricted to transcriptomic and global protein-abundance data.

#### Cardiac Gene2Phenotype and ClinGen analysis

Cardiac gene–disease relationships and pathogenic mechanisms were obtained from the Cardiac Gene2Phenotype^16^ resource and mapped to quantified rat RNA and protein responses using human ortholog annotations. Definitive and strong Gene2Phenotype classifications were considered high confidence, with moderate-confidence relationships also retained. ClinGen dosage-sensitivity annotations^17^ were additionally used to assess evidence for haploinsufficiency. For genes with high or moderate confidence and an interpretable dosage- or activity-related disease mechanism, the latest training time point showing a nominal P ≤0.05 response in either sex or molecular layer was evaluated. Increased abundance for reduced-dosage/loss-of-function mechanisms, or decreased abundance for increased-activity/gain-of-function mechanisms, was classified as potentially compensatory; the opposite relationships were classified as potentially adverse, and discordant responses as mixed. High-potential compensatory classifications required either a monoallelic reduced-dosage mechanism with established ClinGen haploinsufficiency evidence or an established increased-activity mechanism.

#### Alignment of peripheral-tissue responses to cardiac temporal programs

To assess whether cardiac molecular programs were conserved across tissues, cardiac factor-associated features measurable in gastrocnemius skeletal muscle or liver were evaluated in the same animals. For each sex and factor, normalized peripheral-tissue feature abundance was correlated with the corresponding cardiac factor scores using Pearson correlation. P values were adjusted within each tissue, molecular layer, sex, and factor, and associations with |r| ≥0.4 and adjusted P ≤0.05 were retained. Peripheral responses were classified as conserved when association with the corresponding cardiac factor was maintained in the same direction, opposed when the direction was reversed, or different trend when the feature was preferentially associated with a different temporal factor. For pathway-level summaries, classified features were grouped into the major cardiac functional categories defined in the corresponding analyses, and the numbers of conserved, opposed, and different trend responses were summarized within each category.

### Supplementary results

#### Sex-dependent feature detection and molecular response concordance

The number of exercise-responsive cardiac features differed between females and males across molecular layers and training durations (**Supplementary Figure 1A**). At stringent FDR thresholds, more significant features were generally detected in males, particularly for transcriptomic and proteomic responses. Relaxing the statistical threshold reduced this imbalance in some comparisons, although differences remained dependent on molecular layer and training duration. To determine whether limited overlap between significant feature lists reflected discordant molecular regulation, we compared female and male training-effect estimates among features showing nominal evidence of responsiveness (P ≤ 0.01 in at least one sex) at each timepoint (**Supplementary Figure 1B**). Effect-size correlations and directional agreement varied across molecular layers. Transcriptomic directional concordance ranged from 69.2% to 83.7%, whereas proteomic concordance increased from approximately 73% during early training to 88.0%–88.7% at 4–8 weeks. Acetylation showed a similar increase at later timepoints (86.1%–88.1%), while phosphosite responses exhibited relatively high directional agreement throughout training (76.2%– 82.5%). Metabolites also showed substantial concordance, reaching 90.8% at 8 weeks. We next examined whether differences in statistical detection were associated with response magnitude or precision (**Supplementary Figure 1C–D**). Among nominally responsive features, female protein effect estimates were approximately 15% and 28% smaller in median absolute magnitude than male estimates at 4 and 8 weeks, respectively, despite comparable standard errors. Transcriptomic effect magnitudes were more similar between sexes at these later timepoints, with standard error ratios also close to unity. Metabolites showed a more pronounced reduction in estimated female response magnitude, accompanied by larger standard errors, particularly at 8 weeks.

Together, these findings suggest that limited overlap between statistically significant feature lists may underestimate the extent of shared directional regulation between sexes. Female and male cardiac responses may therefore be more concordant than significance-based comparisons suggest, particularly at later training stages, despite smaller estimated response magnitudes in females across several molecular layers.

#### Multi-omic model definition

For multi-omic integration, features were selected based on nominal evidence of training responsiveness (P ≤ 0.01 in at least one sex-specific timewise contrast or overall-training test), with additional restrictions for epigenetic regions associated with genes showing RNA or protein responses (see Methods). The resulting dataset comprised 13,790 features: 2,457 transcripts, 1,996 DNA methylation sites, 447 chromatin accessibility peaks, 2,091 proteins, 897 acetylation sites, 5,324 phosphosites, 52 ubiquitination sites, and 526 metabolites.

To determine model dimensionality, MOFA+/MEFISTO models containing 2–15 factors were evaluated using 10 independent random seeds per configuration (**Supplementary Figure 3**). Increasing the number of factors improved the mean variance explained across molecular layers (**Supplementary Figure 3A**) and increased the number explaining substantial molecular variation (≥2% variance in at least two molecular views and ≥5% cumulatively, **Supplementary Figure 3B**). However, higher-dimensional models also produced more factors dominated by a single molecular layer (**Supplementary Figure 3C**). Seven factors were therefore retained, capturing >20% mean variance across molecular layers in both sexes while limiting omic-specific factors.

#### Extended functional characterization of multi-omic temporal programs

To further characterize the biological processes underlying these temporal programs, pathway enrichment was examined separately for positively and negatively weighted features within each molecular layer. Within the sustained Factor 2 program, complement and coagulation pathways were strongly enriched among negatively weighted proteins (P = 2.27 × 10⁻¹¹). Muscle-cell cytoskeleton pathways were enriched among positively weighted phosphosites (P = 7.39 × 10⁻¹⁰) and proteins (P = 0.014), but negatively weighted transcripts (P = 0.0049), potentially reflecting distinct regulation at the transcript, protein, and phosphorylation levels during sustained structural remodeling. Additional transcriptomic enrichments involved integrin signaling and cardiac conduction. Metabolic processes were also represented, with TCA-cycle and fatty-acid oxidation pathways enriched among acetylation sites and concordant increases in Eno3 transcript and protein abundance in both sexes, supporting a shared, sustained component of cardiac metabolic remodeling.

Sex-consistent transient programs involved complementary metabolic, translational, and structural processes. Factor 4 combined enrichment of PPAR signaling (P = 1.50 × 10⁻⁷) and fatty-acid oxidation (P = 1.67 × 10⁻⁶) among positively weighted transcripts with ribosomal pathways among negatively weighted transcripts (P = 1.07 × 10⁻⁸), consistent with early changes in substrate handling and translational regulation. Factor 3 was associated with muscle-structure development among negatively weighted acetylation features (P = 0.0098). Together, these findings suggest an early response shared across sexes, combining metabolic and translational adjustments with more selective structural remodeling.

Programs with sex-specific temporal trajectories showed prominent mitochondrial and structural associations. Factor 5 exhibited a transient increase in females but a modest decrease in males. Its negatively weighted acetylation sites were enriched for aerobic respiration and respiratory electron transport (P = 2.57 × 10⁻⁴), TCA-cycle processes, and fatty-acid oxidation, while negatively weighted transcripts were associated with translational and muscle-contractile pathways. These opposing trajectories suggest contrasting regulation of shared molecular features between sexes, without necessarily implying opposite changes in pathway activity. In contrast, Factor 6 followed a similar direction of response in both sexes, emerging earlier in males but delayed until 4 weeks in females. Its extensive proteomic enrichments encompassed branched-chain amino-acid degradation (P = 1.51 × 10⁻²⁶), aerobic respiration and electron transport (P = 3.30 × 10⁻²⁵), TCA-cycle metabolism, and fatty-acid oxidation. Factor 1 captured an additional early female response, with reduced chromatin accessibility at 1 week and enrichment of associated ATAC-seq peaks for glucagon signaling and cellular stress pathways. Although ATAC-seq responses to training were relatively limited, these early changes may provide indirect support for regulatory differences associated with delayed metabolic remodeling in females. Finally, Factor 7 combined enrichments also represented in transient and sustained programs, including positively weighted transcripts associated with PPAR signaling (P = 4.50 × 10⁻⁸) and oxidative metabolism, and negatively weighted transcripts associated with cytoskeletal (P = 6.40 × 10⁻⁷), integrin-signaling, and extracellular-matrix pathways, reflecting the sex-specific sustained or transient programs.

#### Cross-tissue comparisons reveal limited structural conservation but broader metabolic correspondence

To determine whether cardiac remodeling was shared with other exercise-responsive tissues, we compared heart with gastrocnemius skeletal muscle (SKMGN) from the same animals across multiple molecular layers and training timepoints. Direct comparisons revealed limited overlap of exercise-responsive features between tissues at matched timepoints (**Supplementary Figure 6A**). We therefore examined whether molecular features defining cardiac temporal programs followed similar trajectories in skeletal muscle (**Supplementary Figure 6B-C, Supplementary Table 6**). Significant temporal alignment was detected in 1,513 of 11,899 evaluations of heart-selected features (12.7%). Among these, only 300 (19.8%) reproduced the corresponding cardiac temporal pattern, whereas 1,063 (70.3%) followed different temporal patterns and 150 (9.9%) showed opposing responses. Thus, even when molecular features responded in both tissues, their temporal regulation was rarely conserved.

To determine whether cross-tissue correspondence differed across biological functions, we grouped the aligned molecular features according to the functional categories identified in the cardiac multi-omic analyses (**Supplementary Figure 6E**). Cardiac structure/function and cardiomyopathy-related features showed particularly limited conservation of temporal trajectories, with most aligned features following different or opposing patterns in SKMGN. FHL1, involved in sarcomeric organization and mechanical stress sensing, provided a conserved example, increasing late in both heart and SKMGN in females and males (**Supplementary Figure 6D**). In contrast, PDLIM3 increased progressively in heart but showed an early transient decrease in female SKMGN, whereas XIRP2 increased in heart while decreasing early in SKMGN in both sexes. These findings indicate that intracellular structural remodeling was largely heart-specific in its temporal organization. ECM remodeling represented a notable exception, with reductions in ECM-associated features largely shared between heart and SKMGN in males (**Supplementary Figure 6F**).

In contrast, metabolic processes showed much broader cross-tissue correspondence, particularly mitochondrial energy and lipid/fuel metabolism across proteomic and metabolomic layers (**Supplementary Figure 7**). However, most aligned features followed different rather than conserved temporal trajectories, indicating that similar metabolic processes were engaged with distinct timing in heart and SKMGN. Consistent with this distinction, skeletal muscle showed more sustained and progressive remodeling of mitochondrial fatty-acid oxidation, TCA-cycle, and oxidative-phosphorylation pathways, compared with earlier or intermediate responses in heart. To distinguish metabolic responses shared between striated muscles from more widespread adaptation, we also examined the liver tissue from the same animals (**Supplementary Figure 7**). Hepatic responses showed substantially less overlap with cardiac molecular changes at matched timepoints and did not reproduce the coordinated mitochondrial remodeling observed in heart and SKMGN, despite changes in individual metabolic features (**Supplementary Figure 6A**). In contrast, selected lipid-associated responses showed broader cross-tissue correspondence. The lipoprotein-associated proteins APOC1, APOC2, APOC3, and APOE decreased progressively in heart, with similar reductions in SKMGN and liver in females (**Supplementary Figure 8A**). Cardiac triglycerides reached their greatest reduction at 4 weeks in females but continued to decline through 8 weeks in males, following a similar sex-dependent pattern in liver (**Supplementary Figure 8B**). These findings indicate that selected aspects of lipid transport and storage were shared across tissues despite more limited correspondence in mitochondrial remodeling. Together, these comparisons highlight the tissue specificity of cardiac structural remodeling, broader but temporally distinct metabolic responses in skeletal muscle, and selective conservation of lipid-associated changes in liver.

#### Additional temporal and regulatory features of cardiac fuel remodeling

Beyond the representative proteins highlighted in the main Results, additional metabolic features provided further resolution of the temporal organization of cardiac lipid remodeling (**Supplementary Figure 7**). Early training involved increased proteins associated with peroxisomal fatty-acid metabolism (ACOX1, PHYH) in both sexes. MLYCD protein also increased at 2–4 weeks, potentially influencing mitochondrial fatty-acid entry by degrading malonyl-CoA, an inhibitor of CPT1. A targeted acyl-CoA feature annotated as malonyl-CoA/hydroxybutyryl-CoA decreased throughout training in males and at 1 and 8 weeks in females, although its unresolved annotation prevented confirmation of reduced malonyl-CoA abundance. In males, CPT2, involved in mitochondrial fatty-acid transport, increased at 2 weeks before decreasing at 8 weeks, highlighting transient remodeling of the carnitine shuttle. These changes were accompanied by reductions in selected acylcarnitines and subsequent increases in long- and very-long-chain fatty acids, the latter preceding the greatest reduction in cardiac triglycerides at 8 weeks. Lipid-droplet regulation also differed between sexes, with *Plin5* transcript abundance decreasing in females but increasing transiently in males before declining by 8 weeks.

Early female carbohydrate remodeling involved additional changes in pentose-phosphate– associated metabolites at 1 week, with increased sedoheptulose-7-phosphate and gluconate but decreased erythrose-4-phosphate (**Figure 5**). Together with the reductions in dihydroxyacetone phosphate and glycerol-3-phosphate described in the main Results, these responses indicated early reorganization of carbohydrate-derived metabolite pools linked to glycerolipid synthesis and redox metabolism, although metabolite abundance does not establish pathway flux. Exploratory ATAC-seq profiles also suggested reduced accessibility at selected glycolytic, pyruvate-metabolic, and mitochondrial loci at 1 week in females, although most individual changes were not statistically significant (**Supplementary Figure 8C**).

Beyond the representative sites described in the main Results, phosphorylation and acetylation involved proteins regulating lipid handling (PLIN5, MLYCD), mitochondrial β-oxidation (HADHA, HADHB), and the TCA cycle (CS, ACO2, IDH2, MDH2; **Figure 5B**). MDH2 exhibited distinct temporal patterns across K239, S326, and K307, highlighting site-specific regulation within individual metabolic enzymes. These modifications provided an additional regulatory dimension to cardiac metabolic remodeling, although their functional consequences cannot be inferred directly from PTM abundance.

Finally, the disease-opposing metabolic response extended beyond the lipoprotein-associated changes highlighted in the main Results to include intracellular substrate metabolism (**Supplementary Figure 8B**). *Eno3* increased throughout training in both sexes, whereas SLC2A4 and HK2, involved in glucose handling, and PNPLA2, involved in lipid mobilization, showed more prominent increases in males. Additional responses involving GPD1 and *Perm1* further connected this disease-opposing signature to carbohydrate metabolism and mitochondrial oxidative regulation.

### Supplementary figures legend

**Supplementary Figure 1. Sex differences in cardiac molecular response detection, concordance, magnitude, and precision during endurance training.**

**A)** Number of exercise-responsive molecular features detected in females and males across progressively relaxed statistical thresholds. Differential features were identified separately by sex at each training timepoint relative to same-sex sedentary controls and summarized across five molecular classes: transcriptomic, epigenetic (DNA methylation and chromatin accessibility), proteomic, post-translational modification (PTM; acetylation, phosphorylation, and ubiquitination), and metabolomic. Lines indicate the number of significant features in males (blue), females (pink), and shared between sexes at the same timepoint (gray).

**B)** Female–male concordance of training-effect estimates across molecular layers and timepoints. Features with nominal p-value≤0.01 in at least one sex were selected independently at each timepoint, and the same selected features were evaluated in both sexes. Heatmap color represents the Spearman correlation (ρ) between female and male effect estimates, and cell values indicate the percentage of features showing concordant directions of change.

**C)** Relative magnitude of female and male training effects among the same nominally responsive features defined in B. Cell values represent the ratio of median absolute log2(fold changes) in females versus males, calculated separately for each molecular layer and timepoint. Ratios below 1 indicate smaller estimated response magnitudes in females, whereas ratios above 1 indicate larger magnitudes.

**D)** Female-to-male standard error ratios among the same selected features. Cell values represent the median of feature-wise female/male standard error ratios for each molecular layer and timepoint. Values above 1 indicate larger standard errors in females, whereas values below 1 indicate larger standard errors in males.

**Supplementary Figure 2. Cross-omic clustering of enriched biological processes reveals recurrent functional programs across training.**

**A)** Multi-omic clustering of significantly enriched Gene Ontology Biological Process (GO:BP) terms. Significant enrichment results from individual molecular layers, sexes, and training timepoints were integrated, and pairwise similarity between enriched GO terms was calculated by combining gene-set overlap and GO semantic similarity. The resulting similarity matrix was hierarchically clustered using average linkage and a distance of 1−similarity; clusters were defined by cutting the dendrogram at a similarity threshold of τ=0.10. Heatmap color indicates pairwise term similarity. Annotations indicate the molecular layer contributing each enriched term and the manually curated biological category assigned to each cluster; smaller clusters (n terms ≤10) are grouped as Other clusters. The resulting clusters comprised terms contributed by multiple molecular layers and resolved recurrent functional themes.

**B)** Temporal and sex-specific enrichment patterns underlying the curated biological categories in A. Significant GO:BP terms (FDR≤0.05) are shown across molecular layers, training timepoints, and sexes. When multiple significant terms from the same category occurred for a given molecular layer, timepoint, and sex, the term with the lowest FDR was displayed. Symbol size represents enrichment significance (−log10[FDR]), outline color indicates sex (female, pink; male, blue), and symbol orientation indicates normalized enrichment score (NES) direction (upward, positive; downward, negative). For transcript and protein abundance, NES direction reflects enrichment toward positively or negatively changing features; for epigenetic and post-translational modification layers, NES direction indicates the direction of change in the corresponding sites or features and does not necessarily imply increased or decreased gene or protein abundance.

**Supplementary Figure 3. Selection of the number of factors for multi-omic integration.**

MOFA models were trained across a range of requested factor numbers (K=2–15) using multiple random seeds; boxplots summarize variation across model runs, and the dotted vertical line indicates the number of selected factors.

**A)** Variance explained (R²) across molecular views for female and male models. For each model, variance explained by all factors was summed within each molecular view and then averaged across views.

**B)** Number of active factors in female and male models and number active in both sexes. A factor was considered active within a sex if it explained ≥2% variance in at least two molecular views and ≥5% total variance across views, or ≥5% variance in a single view.

**C)** Number of single-view factors, defined as active factors predominantly represented by a single molecular view rather than across multiple views. K=7 was selected as a parsimonious solution that captured >20% mean variance across molecular views in both sexes while preceding the greater emergence of active and single-view factors at higher K.

**Supplementary Figure 4. Selected MOFA-associated features recapitulate factor temporal trajectories.**

Sex-specific temporal trajectories of features associated with each MOFA factor across the 8-week training course. Features were selected separately in females and males based on the correlation between their normalized abundance and the corresponding MOFA factor scores (Pearson |r|≥0.4 and FDR≤0.05) and stratified according to positive (+) or negative (−) MOFA weights. Normalized abundances for each feature were converted to Z-scores across samples and then averaged within each training timepoint. Individual feature trajectories are shown in gray, with greater opacity indicating closer agreement with the consensus trajectory. Colored lines represent the mean trajectory across selected features and shaded ribbons indicate ±SD; solid and dashed lines denote positively and negatively weighted features, respectively. The number of features contributing to each trajectory is indicated within each panel. Female and male trajectories are shown in pink and blue, respectively.

**Supplementary Figure 5. Rat ortholog coverage of CVD Atlas disease signatures.**

Bars indicate the fraction of unique CVD Atlas disease-associated genes or proteins that could be mapped to rat orthologs, stratified by disease and omic layer. Transcriptomic counts were substantially higher than proteomic counts, reflecting the greater representation of transcriptomic datasets in the cardiac CVD Atlas subset (24 transcriptomic versus 5 proteomic datasets; 15,337 versus 534 disease-associated gene/protein instances before rat mapping).

**Supplementary Figure 6. Cross-tissue comparison of exercise-responsive cardiac features and temporal programs. A)** Heatmap showing overlap of exercise-responsive features between heart and peripheral tissues at each training timepoint by sex (p-value ≤0.01). Numbers indicate the count of differentially affected features shared with heart, matched by sex, timepoint, and molecular layer. Cell shading indicates the percentage of differentially affected features in each peripheral tissue represented by these shared features. Gray cells indicate unavailable assays. RRBS, ATAC-seq, and ubiquitination were excluded. SKMGN: gastrocnemius skeletal muscle.

**B)** Alluvial plot showing temporal alignment of heart-selected molecular features in gastrocnemius (SKMGN) skeletal muscle in females and males. Features were classified as conserved, different trend, or opposed according to their best significant alignment with cardiac factor trajectories (adjusted p-value ≤0.05, |r|≥0.4). Percentages indicate the proportion of evaluable heart-selected features with significant skeletal muscle alignment. Right-hand flows show the molecular-layer composition of aligned features. RRBS, ATAC-seq, and ubiquitination were excluded.

**C)** Temporal trajectories of skeletal muscle features correlated with cardiac factor programs in SKMGN. Features with significant positive or negative correlations with the corresponding cardiac factor trajectory (adjusted P≤0.05, |r|≥0.4) were grouped by sex and cardiac temporal pattern. Thin gray lines show individual feature trajectories based on normalized expression z-scores; colored lines and ribbons show the median and interquartile range for positively (red) and negatively (blue) correlated features, respectively. Black dashed lines indicate the corresponding cardiac factor trajectory. Annotated n values indicate the number of correlated features in each group.

**D)** Representative cardiac–skeletal muscle temporal responses of the structural proteins Fhl1, Pdlim3, and Xirp2. Normalized log-transformed abundance is expressed relative to the median sedentary value within each tissue and sex. Boxplots show the median and interquartile range. Statistical significance reflects comparison of each exercise timepoint with sedentary controls within the same tissue and sex. p-value *≤0.01.

F) Number of features associated with each major cardiac-enriched biological function across skeletal muscle tissues and available molecular layers, stratified by temporal alignment class. Counts represent unique genes for transcript, protein, PTM (acetylation and phosphorylation), and metabolite features.

**F)** Dumbbell plot showing 8-week log2(FC) of selected genes and proteins associated with cardiac-enriched extracellular matrix functions in HEART and SKMGN, separately by sex. Values are relative to sedentary controls. Filled symbols indicate significant responses (raw p-value ≤0.01), whereas open symbols indicate nonsignificant responses. Red: HEART; blue: SKMGN.

**Supplementary Figure 7. Cross-tissue temporal remodeling of metabolic pathways during endurance training.**

Heatmaps represent exercise-induced changes in representative transcript and protein features across heart, gastrocnemius skeletal muscle (SKMGN) and liver. Metabolic programs are organized into fuel handling and entry (fatty-acid handling, peroxisomal lipid metabolism, mitochondrial fatty-acid oxidation, and pyruvate routing/PDH regulation), mitochondrial carbon oxidation (BCAA/amino-acid input and TCA cycle), and oxidative capacity and mitochondrial support (ETC/OXPHOS, mitochondrial gene expression/maintenance, and mitochondrial cofactor/redox metabolism). Heatmap values indicate log₂FC relative to sex-matched sedentary controls at each training timepoint. Asterisks indicate nominal p-value≤0.01 versus sedentary controls. Side annotations denote molecular layer, tissue, sex, and training duration.

**Supplementary Figure 8. Regulatory and cross-tissue features of sex-specific metabolic remodeling with endurance training. A)** Cross-tissue comparison of selected metabolic features contributing to the disease-opposing cardiac response. Heatmaps show log2 fold changes relative to sedentary controls across training timepoints in HEART, gastrocnemius skeletal muscle (SKMGN), and LIVER, separately by sex and molecular layer. Asterisks indicate significant changes relative to same-sex sedentary controls (nominal P≤0.01). The side annotation indicates biological function, and the cardiac-disease annotation identifies features previously classified as oppositely regulated between exercise and human cardiovascular disease signatures.

**B)** Temporal profiles of triglyceride abundance in heart and liver. All measured triglyceride features were standardized separately within each tissue and sex across sedentary and training groups (132 features in heart; 151 in liver). For each animal, the median z-score across triglyceride features was calculated to derive a relative triglyceride-class abundance score. Lines connect group medians and shading indicates the interquartile range across animals at each timepoint. Female and male trajectories are shown in pink and blue, respectively.

**C)** Chromatin accessibility at selected loci associated with cardiac glycolytic, pyruvate-metabolic, and mitochondrial substrate-handling genes. Boxplots show normalized ATAC-seq accessibility for peaks associated with the indicated genes across training timepoints and sexes. Boxes indicate the median and interquartile range. Gray curves show sex-specific LOESS trajectories fitted to individual animals, with shading indicating the 50% confidence band. Asterisks indicate differential accessibility relative to same-sex sedentary controls from timewise DESeq2 analyses (p-value *≤0.05, **≤0.01).

**Supplementary Figure 9. Proteostatic, redox, immune, and hemostatic remodeling across tissues during endurance training.**

Heatmaps show training-induced log₂ fold changes across heart, gastrocnemius skeletal muscle (SKMGN), and liver for (**A**) proteostasis, mitochondrial redox, and cytoprotective proteins, (**B**) immune/stress-related transcripts, and (**C**) complement, contact/kinin, coagulation/fibrinolysis, and plasma protease-regulatory proteins. Female and male responses are shown at 1, 2, 4, and 8 weeks relative to sex-matched sedentary controls; cardiac-disease annotations indicate similar, opposed, mixed, or no overlap with cardiovascular-disease signatures. Asterisks indicate nominal P ≤ 0.01.

### Supplementary tables legend

**Supplementary Table 1. Cardiac MOFA sample factor scores.**

Sample-level scores for the seven cardiac factors, with sample ID, sex, and training timepoint.

**Supplementary Table 2. Sex-specific associations between molecular features and cardiac MOFA factors.**

Each row represents a molecular feature with at least one qualifying association. F, M, and F+M indicate associations meeting both statistical thresholds in females, males, or both sexes, respectively (absolute correlation ≥ 0.40 and FDR ≤ 0.05). F+M does not necessarily indicate concordant correlation directions between sexes. Blank cells indicate no qualifying association. Position denotes amino acid residues for post-translational modifications or genomic intervals for epigenetic features.

**Supplementary Table 3. Significant pathway enrichments for cardiac MOFA factors.**The table reports 380 significant over-representation results from Gene Ontology (GO) Biological Process, Reactome, and KEGG. Enrichment analyses were performed separately by factor, molecular layer, and MOFA weight sign, using the combined features selected in females and males. Each row reports the g:Profiler g:SCS-adjusted P value, number of overlapping genes, and contributing rat genes.

**Supplementary Table 4. Direction of cardiac exercise responses relative to human cardiovascular disease signatures.**

The table summarizes transcript and protein direction calls for 425 genes across 10 cardiovascular disease categories within sustained cardiac programs. Aligned and Opposed indicate changes in the same or opposite direction as the disease signature, respectively. Mixed indicates discordant transcript and protein direction calls for the same gene, disease, and sex. Blank cells indicate no qualifying direction call. Female and male columns are shaded pink and blue, respectively. HCM, hypertrophic cardiomyopathy; DCM, dilated cardiomyopathy; ICM, ischemic cardiomyopathy; HF, heart failure; LVHF, left ventricular heart failure; BiVHF, biventricular heart failure; HFrEF and HFpEF, heart failure with reduced and preserved ejection fraction, respectively; MyoI, myocardial ischemia; CH, compensated hypertrophy.

**Supplementary Table 5. Inherited disease associations and dosage evidence for cardiac genes.**

The table lists 41 Gene2Phenotype (G2P) records covering 29 genes, including the associated disease, molecular mechanism, variant consequence, allelic requirement, and G2P confidence rating. ClinGen haploinsufficiency and triplosensitivity classifications were obtained from the June 7, 2026 dosage sensitivity snapshot. “Not in snapshot” indicates that the gene had no entry in that snapshot.

**Supplementary Table 6. Best matches to cardiac temporal programs in gastrocnemius skeletal muscle.**

Each row reports the best qualifying gastrocnemius skeletal muscle (SKMGN) match for a molecular feature, sex, and cardiac factor (absolute correlation ≥ 0.40 and FDR ≤ 0.05). Conserved, Opposed, and Different trends indicate similar, opposed, or distinct temporal response patterns, respectively. The table includes 198 conserved, 20 opposed, and 527 different-trend matches. Alignment percentages are calculated using all 4,573 eligible assignments, including 3,828 assignments without a qualifying match that are omitted from the table.

## MoTrPAC Study Group

### Consortium Coordination

**Bioinformatics Center - Stanford University, Stanford, CA**

David Amar; Euan Ashley; Pauline Brochet; Pauline Brochet; Sasha Gladkikh; David Jimenez-Morales; Daniel H. Katz; Daniel Seung Kim; Malene E. Lindholm; Samuel Montalvo; Joyce Njoroge; Nikolai G. Vetr; Matthew T. Wheeler;

**Data Management, Analysis, and Quality Control Center - Wake Forest University School of Medicine, Winston-Salem, NC**

Jerry Barnes; Brandon Bukas; Haiying Chen; Shyh-Huei Chen; Josh Cherian; Shannon S. Emilson; Catherine Gervais; Leora Henkin; Fang-Chi Hsu; Guido Mascia; Ryan McGinnis; Michael E. Miller; John Nichols; David Popoli; Eric W. Reynolds; Jeremy Rogers; Scott Rushing; Santiago Saldana; Courtney Simmons; Cynthia L. Stowe; Jennifer W. Talton; Michael P. Walkup;

**Exercise Intervention Core - Wake Forest University, Winston-Salem, NC**

Jason Fanning; W. Jack Rejeski;

**National Institutes of Health, Bethesda, MD**

Amanda Boyce; Emily Carifi; Jana Eisenstein; Jerome L. Fleg; Lyndon Joseph; Padma Maruvada; Concepcion R. Nierras; George Papanicolaou; John P. Williams; Ashley Xia;

### Chemical Analysis Sites

**Broad Institute of MIT & Harvard (Carr Lab), Cambridge, MA**

Steven Carr; Pierre M. Jean Beltran; Hasmik Keshishian; Charles C. Mundorff;

**Broad (Newgard Lab) - Duke University, Durham, NC**

Danhong Lu; Michael Muehlbauer; Christopher B. Newgard; **Georgia Institute of Technology, Atlanta, GA**

Carter Asef; Facundo M. Fernández; David A. Gaul; Samuel G. Moore;

**Icahn School of Medicine at Mount Sinai, New York City, NY**

Mary Anne S. Amper; Yongchao Ge; Venugopalan D. Nair; German Nudelman; Hanna Pincas; Stuart C. Sealfon; Gregory R. Smith; Gregory R. Smith; Mital Vasoya; Martin J. Walsh; Martin J. Walsh; Elena Zaslavsky;

**Pacific Northwest National Laboratory, Richland, WA**

Joshua N. Adkins; Marina A. Gritsenko; Chelsea Hutchinson-Bunch; Gina M. Many; Wei-Jun Qian; James A. Sanford;

**Stanford University, Stanford, CA**

Abdalla Ahmed; Jose Juan Almagro Armenteros; Nasim Bararpour; Clarisa Chavez; Nicole R. Gay; Krista M. Hennig; Chia-Jui Hung; Christopher A. Jin; Curtis McGinity; Stephen B Montgomery; Daniel Nachun; Kevin S. Smith; Michael P Snyder; Si Wu; Yilin Xie; Navid Zebarjadi; Bingqing Zhao;

**Preclinical Animal Study Sites**

**Joslin Diabetes Center, Boston, MA**

Laurie J. Goodyear; Michael F. Hirshman;

**University of Florida, Gainesville, FL**

Karyn A. Esser;

**University of Iowa, Iowa City, IA**

Dam Bae; Sue C. Bodine; Simon Schenk;

**Virginia Polytechnic Institute and State University, Charlottesville, VA**

Matthew Wolf; Zhen Yan;

### Clinical Sites

**Ball State University, Muncie, IN**

Bridget Lester; Scott Trappe; Todd A. Trappe;

**The University of Alabama at Birmingham - Birmingham, AL**

Marcas M. Bamman; Thomas W. Buford; Gary R. Cutter;

**Duke University, Durham, NC**

William E. Kraus;

**University of California, Irvine, CA**

Dan M. Cooper; Fadia Haddad; Shlomit Radom-Aizik;

**University of Colorado, Denver, CO**

Bryan C. Bergman; Zachary S. Clayton; Wendy M. Kohrt; Edward L. Melanson; Kerrie L. Moreau;

**University of Texas Health Science Center, San Antonio, TX**

Blake B. Rasmussen;

## MoTrPAC Study Group Affiliations

**Ball State University, Muncie, IN**

Bridget Lester, Scott Trappe, Todd A. Trappe

**Broad Institute of MIT and Harvard, Cambridge, MA**

Steven Carr, Pierre M. Jean Beltran, Charles C. Mundorff

**Data Management, Analysis, and Quality Control Center, Wake Forest University School of Medicine, Winston-Salem, NC**

Eric W. Reynolds

**Department of Genetics, Stanford University, Stanford, CA**

Euan Ashley, Jose Juan Almagro Armenteros, Nasim Bararpour, Clarisa Chavez, Nicole R. Gay, Christopher A. Jin, Michael P Snyder, Bingqing Zhao

**Department of Neurology, Icahn School of Medicine at Mount Sinai, New York, NY**

Stuart C. Sealfon

**University of Iowa, Iowa City, IA**

Simon Schenk

**Department of Pathology, Stanford University, Stanford, CA**

Euan Ashley, Kevin S. Smith

**Departments of Pathology and Medicine, Stanford University, Stanford, CA**

Nikolai G. Vetr

**Duke Molecular Physiology Institute, Division of Endocrinology, Metabolism and Nutrition, Department of Medicine, Duke University School of Medicine, Durham, NC**

Michael Muehlbauer

**Department of Pharmacology & Cancer Biology and Department of Medicine, Duke University Medical Center, Durham, NC**

Danhong Lu, Christopher B. Newgard

**Duke Molecular Physiology Institute, Duke University, Durham, NC**

Danhong Lu, Christopher B. Newgard

**Georgia Institute of Technology, Atlanta, GA**

Facundo M. Fernández

**Icahn School of Medicine at Mount Sinai, New York, NY**

Mary Anne S. Amper, Yongchao Ge, Venugopalan D. Nair, German Nudelman, Hanna Pincas, Mital Vasoya, Elena Zaslavsky

**Joslin Diabetes Center, Harvard Medical School, Boston, MA**

Laurie J. Goodyear

**Pacific Northwest National Laboratory, Oregon Health & Science University, and The University of Texas at El Paso**

Joshua N. Adkins

**Section on Integrative Physiology and Metabolism, Joslin Diabetes Center, Boston, MA**

Michael F. Hirshman

**Stanford University, Stanford, CA**

Abdalla Ahmed, Krista M. Hennig, Chia-Jui Hung, Curtis McGinity, Stephen B Montgomery, Navid Zebarjadi

**Gray School of Medical Sciences, Tel Aviv University, Tel Aviv, Israel**

David Amar

**University of Alabama at Birmingham, Birmingham, AL**

Marcas M. Bamman

**University of Alabama at Birmingham and Birmingham/Atlanta VA GRECC, Birmingham, AL**

Thomas W. Buford

**University of Alabama at Birmingham School of Public Health, Birmingham, AL**

Gary R. Cutter

**University of California, Irvine, CA**

Dan M. Cooper, Fadia Haddad, Shlomit Radom-Aizik

**University of Colorado Anschutz Medical Campus, Aurora, CO**

Bryan C. Bergman, Zachary S. Clayton, Wendy M. Kohrt, Kerrie L. Moreau

**Division of Endocrinology, Metabolism, and Diabetes, University of Colorado School of Medicine, Anschutz Medical Campus, Aurora, CO**

Edward L. Melanson

**Division of Geriatric Medicine, Department of Medicine, University of Colorado Anschutz Medical Campus, Aurora, CO**

Edward L. Melanson

**University of Texas Health Science Center at San Antonio, San Antonio, TX**

Blake B. Rasmussen

**University of Virginia, Charlottesville, VA**

Matthew Wolf

**Virginia Polytechnic Institute and State University, Charlottesville, VA**

Zhen Yan

## MoTrPAC Study Group Acknowledgements

Carter Asef, Dam Bae, Amanda Boyce, Emily Carifi, Jana Eisenstein, Karyn A. Esser, Jerome L. Fleg, David A. Gaul, Marina A. Gritsenko, Chelsea Hutchinson-Bunch, Lyndon Joseph, Padma Maruvada, Samuel G. Moore, Daniel Nachun, Concepcion R. Nierras, George Papanicolaou, Wei-Jun Qian, James A. Sanford, John P. Williams, Si Wu, Ashley Xia, Yilin Xie

